# Mycobiome analysis in fungal Infected formalin-fixed and paraffin-embedded tissues for identification of pathogenic fungi: A pilot study

**DOI:** 10.1101/2020.04.19.045856

**Authors:** Taebum Lee, Hee Young Na, Kyoung-Mee Kim, Hey Seung Lee, Sung-Hye Park, Ji-Young Choe, Kyoung Chan Choi, Sun-ju Byeon

## Abstract

**Background:** Fungal organisms are frequently observed in surgical pathological diagnosis. In order to more accurately identify fungi in formalin-fixed and paraffin-embedded (FFPE) tissues, it is necessary to use genomic information. The purpose of our pilot study is to identify the factors to be considered for the identification of pathogenic fungi using mycobiome analysis in FFPE tissues.

**Methods:** We selected 49 cases in five hospitals. In each case, FFPE tissue was cut into 50 µm and DNA was extracted. Multiplex PCR with four primers (ITS1, ITS2, ITS3 and ITS4) was performed. Multiplex sequencing was performed using MinION device according to the manufacturer’s protocol. Sequences of each case were searched using BLASTN with an ITS database from NCBI RefSeq Targeted Loci Project with default parameter.

**Results:** A total of 2,526 DNA nucleotides were sequenced. We were able to identify 342 fungal nucleotides in 24 (49.0%, 24/49) cases. The median value of the detected fungal DNA per case was 3 (1Q: 1 and 3Q: 14.25). The 215 (62.87%) fungal DNA contained the entire region of ITS1 or ITS2. The remaining 127 fungal DNAs were identified as fungi using partial sequence of ITS1, ITS2, 5.8S, LSU or SSU.

**Conclusion:** In conclusion, we have identified the possibility of finding pathogenic fungi through mycobiome analysis in fungal infected FFPE tissues using nanopore sequencing method. However, we have also found several limitations to be solved for further studies. If we develop a method to characterize pathogenic fungi in FFPE tissues in a follow-up study, we think it will help patients to use appropriate antifungal agents.

## Introduction

Fungal organisms are frequently observed in surgical pathological diagnosis. Recent developments in technology have enabled the identification of fungi using a variety of methods.^1^ These novel methods are based on analyzing fungal genomes or proteomes using fresh tissues.^2,3^ However, these methods are difficult to apply in formalin-fixed and paraffin-embedded (FFPE) tissues used in surgical pathology. Sanger sequencing and IHC staining have been used in FFPE tissue, but most medical institutions only identify fungi based on morphological findings.^4-9^ The identification of fungi only by morphological findings can lead to inadequate treatment for patients due to misdiagnosis, which can often result in fatal consequences.^10^ Because of the different antifungal agents preferentially used at the initial infection stage depending on the fungus, it is necessary to identify the exact fungi present through testing methods in addition to the morphological findings.^11^

In order to more accurately identify fungi in FFPE tissues, it is essential to use genomic information. DNA markers that could be used to identify fungi include the internal transcribed spacer (ITS) region, small subunit (nrSSU-18S), large subunit (nrLSU-26S or 28S), elongation factor 1-alpha (EF1α), and the largest (RPB1) and second largest (RPB2) subunits of RNA polymerase.^1,2^ Among these markers, ITS could be relatively easily and effectively used for fungal identification, and the database of the ITS regions of fungi is available.^12-14^ In general, sequencing equipment is classified into first- (Sanger sequencing), second- (massively parallel sequencing), and third-generation (real-time and single molecule sequencing) equipment according to key analytical methods.^15^ Third generation sequencing technology is characterized by direct sequencing of nucleotides without PCR amplification. Oxford Nanopore Technology (ONT) introduced several sequencing equipment using the nanopore sequencing technology, which measures the change in current that occurs when a nucleotide sequence passes through a narrow channel.^16^ Since there is an advantage of identifying each DNA sequence without PCR amplification, it is expected that the DNA can be effectively detected in spite of DNA degradation during FFPE tissue preparation and storage.

Extracting only fungal DNA from the fungal infection site of FFPE tissue is very difficult. Therefore, we decided to use the mycobiome analysis method. However, no studies have attempted to analyze mycobiome in FFPE tissue. Therefore, this study was conducted as a pilot study on the development of a method for finding pathogenic fungi using mycobiome analysis in the fungal infected FFPE tissues.

## Materials and Methods

### Sample collection

The cases were extracted from the pathological examination reports of five medical institutions. We first extracted the pathology report that mentioned the presence of fungi. We then selected typical cases for use as positive controls, cases reported to be difficult to differentiate (in briefly, fungi with branched-hyphae, such as *Aspergillus* species and *Mucor* species, are often difficult to distinguish morphologically), and cases with additional information related with fungal identification on pathology report (i.e., identification using culture or sequencing). Through this process, we finally selected 49 cases (case 3-02 and 5-10 were able to confirm the fungal identification results using sequencing). Cases included 21 lungs, 8 paranasal sinuses, 7 gastrointestinal tracts, 5 orbits, 2 skin and mouths, 1 adrenal gland, bone, gum and liver. The average storage period of FFPE tissue was 4.4 years. This study was exempted from obtaining informed consent by Institutional Review Board of Hallym University Dongtan Sacred Heart Hospital (NON2018-005).

### DNA extraction, PCR amplification, Sequencing and Base calling

In each case, FFPE tissue was cut into 50 µm (5 µm × 10) sections and collected in a 2.0 ℳ𝓁 conical tube. DNA was extracted using the ReliaPrep FFPE gDNA Miniprep System (Promega, Madison, Wisconsin, USA) according to the manufacturer’s protocol. The DNA extracted from all cases was loaded onto agarose gel to confirm the presence of DNA. Four universal primers (ITS1, 5’-TCCGTAGGTGAACCTGCGG-3’; ITS2, 5’-GCTGCGTTCTTCATCGATGC-3’; ITS3, 5’-GCATCGATGAAGAACGCAGC-3’; and ITS4, 5’-TCCTCCGCTTATTGATATGC-3’) were ordered from Macrogen (Seoul, Korea) with polyacrylamide gel electrophoresis purification.^17^ Multiplex PCR with four primers was performed as follows: initial denaturation at 95 °C for 5 min, followed by 35 cycles of 94 °C for 30 s, 52 °C for 30 s, and 72 °C for 1 min with a final extension step of 72 °C for 8 min and cooling at 4 °C.^1^ Sequencing was performed using the FLO-MIN106 flow cell (ONT, Oxford Science Park, UK), ligation sequencing kit 1D (SQK-LSK109, ONT) and PCR barcoding expansion 1-12 (EXP-PBC001, ONT) with a MinION (ONT) device in accordance with the manufacturer’s protocol. Multiplex sequencing was performed for 10-12 cases at a time (83-100 ng DNA per sample). The base calling was carried out according to ONT’s recommendations without parameter changes.

### Data analysis

We used BLAST+2.10.0 on a local PC (Ubuntu 19.10) for analysis ^18^. In brief, we downloaded the ITS database file from NCBI RefSeq Targeted Loci Project (last modified Mar 3, 2020) and then converted it using “makeblastdb” of BLAST+2.10.0. This database file contained full or partial ITS sequence of 11,133 genus of fungi. The sequences of *Pneumocystis jirovecii*, which is not found in this database, were downloaded RU7 genomic reference sequences and compared. The sequence of *Actinomyces israelii*, a bacterium that looks morphologically similar to that of a filamentous fungus, was downloaded (*Actinomyces israelii* DSM 43320, whole genome shotgun sequencing project; NZ_JONS00000000.1) and compared. We confirmed the BLASTN search using GRCh38 to determine if the detected DNA corresponds to the human genome.

We converted FASTQ files to FASTA format using “seqtk” (https://github.com/lh3/seqtk). Sequences of each case were searched using BLASTN using ITS database file without default parameter modification. When the sequence matched in the ITS database, the result with the highest bit score value was selected. We selected five results in order of higher bit score when five or more fungal DNAs were detected in one case.

We downloaded the GenBank files of all fungi contained in the database and separated the base sequences of ITS1, 5.8S and ITS2 for each fungus. When fungi were detected, they were classified into three categories (“entire”, “partial”, and “none” match) according to the relationship between the detected fungal DNA and ITS1/ITS2. “Entire” means that all ITS1 or ITS2 base information is used for fungal identification, “partial” means that even if only one sequence information is used, and “none” means that base information other than ITS1 or ITS2 is used.

## Results

A total of 2,526 DNA nucleotides were sequenced. DNA was not seq]]uenced in three cases. Of the remaining 46 cases, the sequenced DNAs of 22 cases were not found in the ITS database. We were able to identify 342 fungal nucleotides in 24 (49.0%, 24/49) cases. The number of DNA sequences identified per case are summarized in Figure 1. The median number of the detected fungal DNA per case was 3 (1Q: 1 and 3Q: 14.25). The clinicopathological information and the number of fungal species detect per case are summarized in Table 1. The remaining cases which fungal DNA was not sequenced and listed in Supplementary Table 1. Detailed information related to the BLAST program is summarized in Supplementary Table 2. The relationship between the detected fungal DNA and the ITS region was summarized in Table 2 and visualized in Figure 2. The 215 (62.87%) fungal DNA contained the entire region of ITS1 or ITS2. The remaining 127 fungal DNAs were identified as fungi using partial sequence of ITS1, ITS2, 5.8S, LSU or SSU.

**Table 1.**
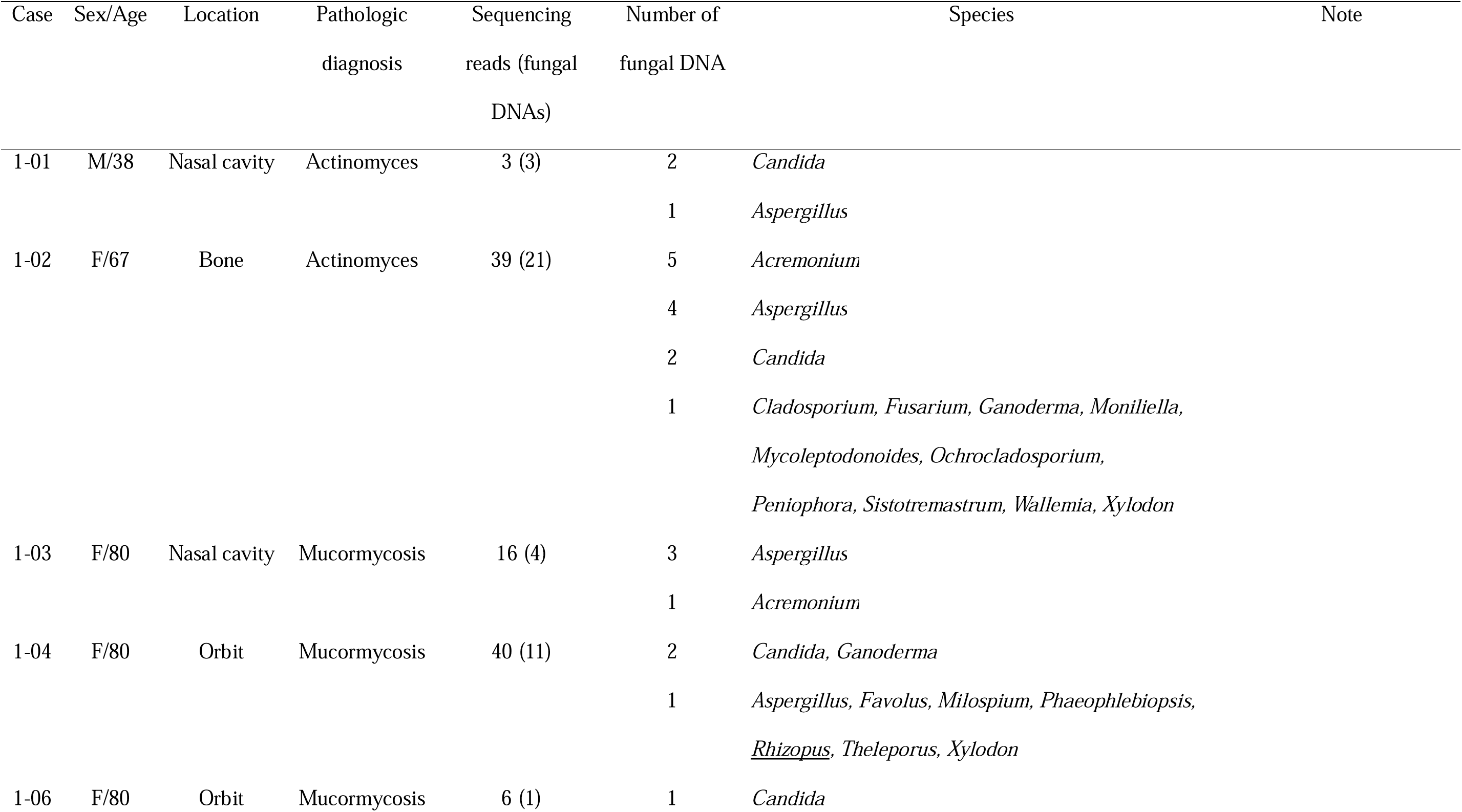
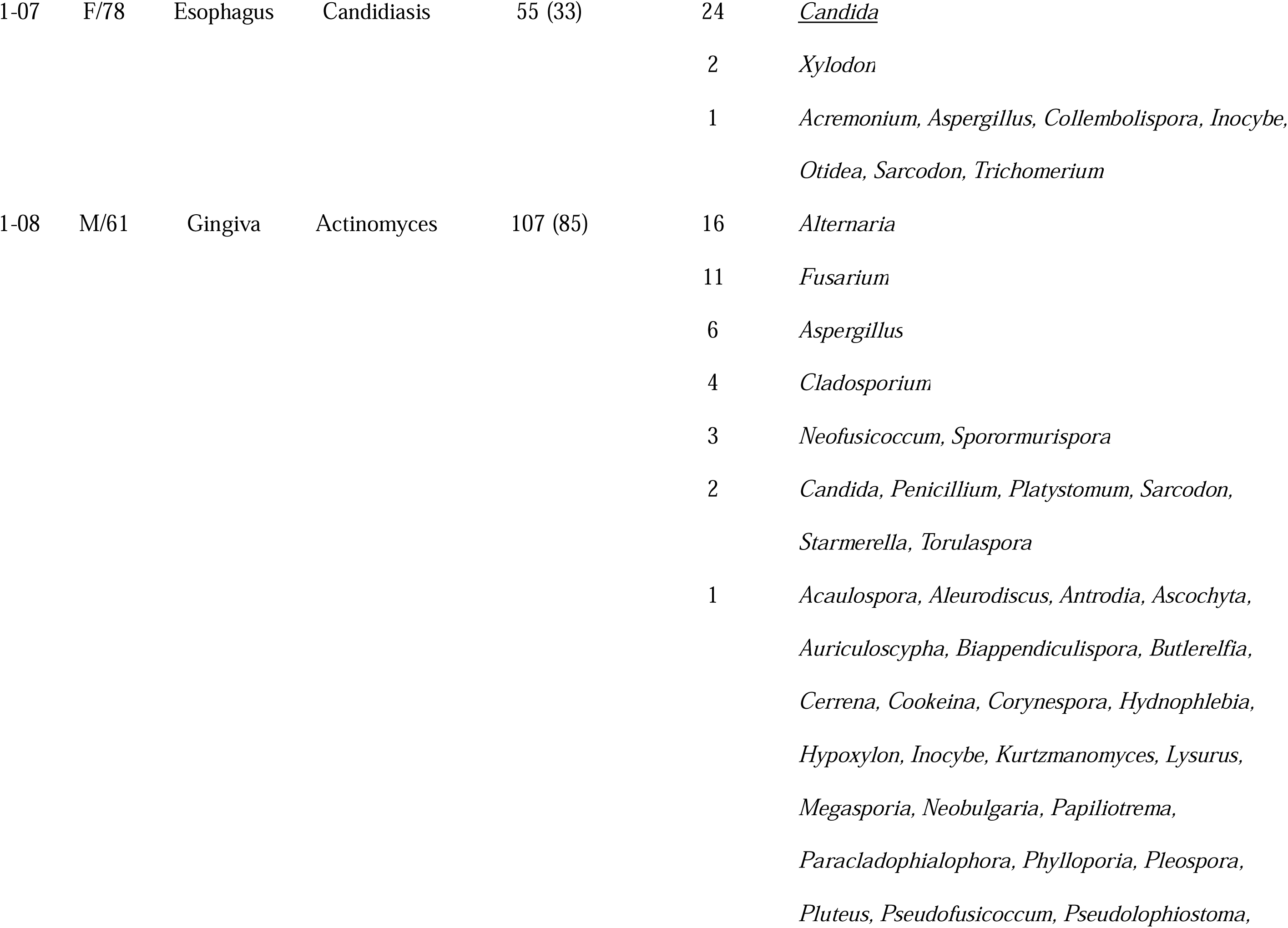
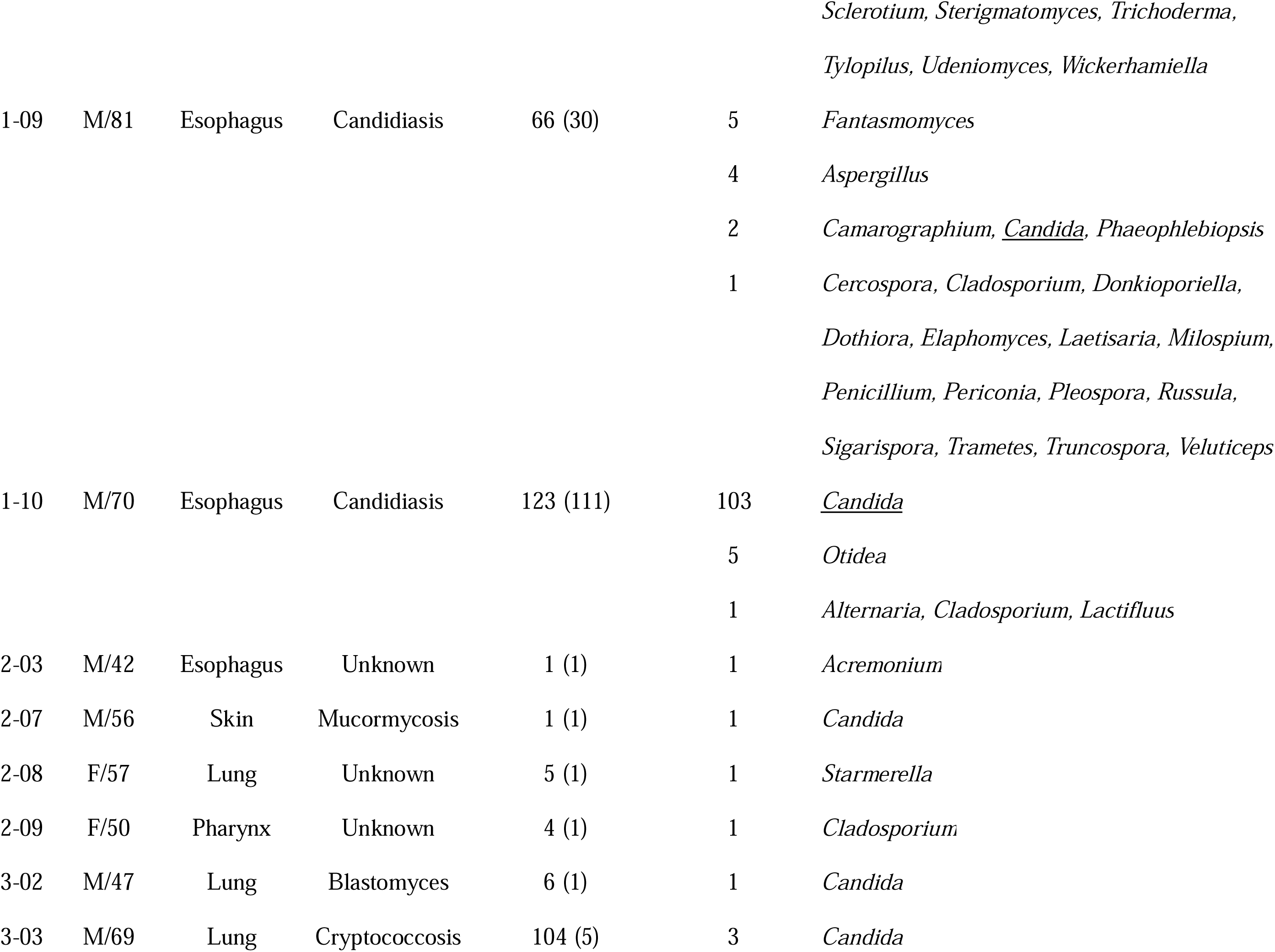
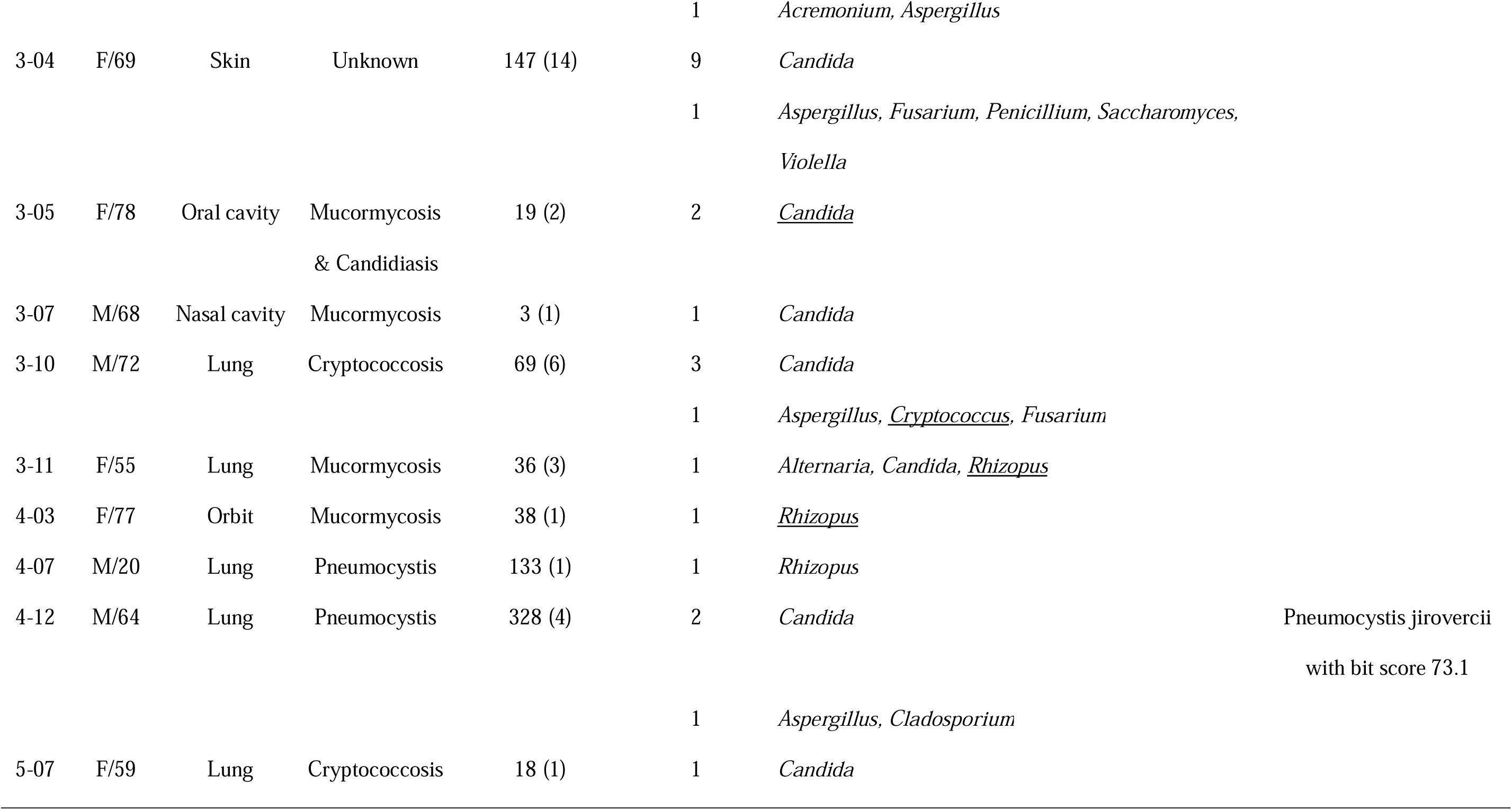
The basic clinicopathological information of fungal DNA confirmed cases (underlined when a fungus identical to pathological diagnosis is detected)

**Table 2.** Classification according to the degree to which the base sequence used to identify fungi matches the ITS1 or ITS2 region (the number in parentheses is percent).

|  |  | ITS2 |  |  |  |
| --- | --- | --- | --- | --- | --- |
|  |  | Entire | Partial | None | Total |
| ITS1 | Entire | 9 (2.63) | 1 (0.29) | 103 (30.12) | 113 (33.04) |
|  | Partial | 4 (1.17) | 1 (0.29) | 58 (16.96) | 63 (18.42) |
|  | None | 98 (28.65) | 51 (14.91) | 17 (4.97) | 166 (48.54) |
|  | Total | 111 (32.46) | 53 (15.50) | 178 (52.05) | 342 (100) |

**Figure 1.**
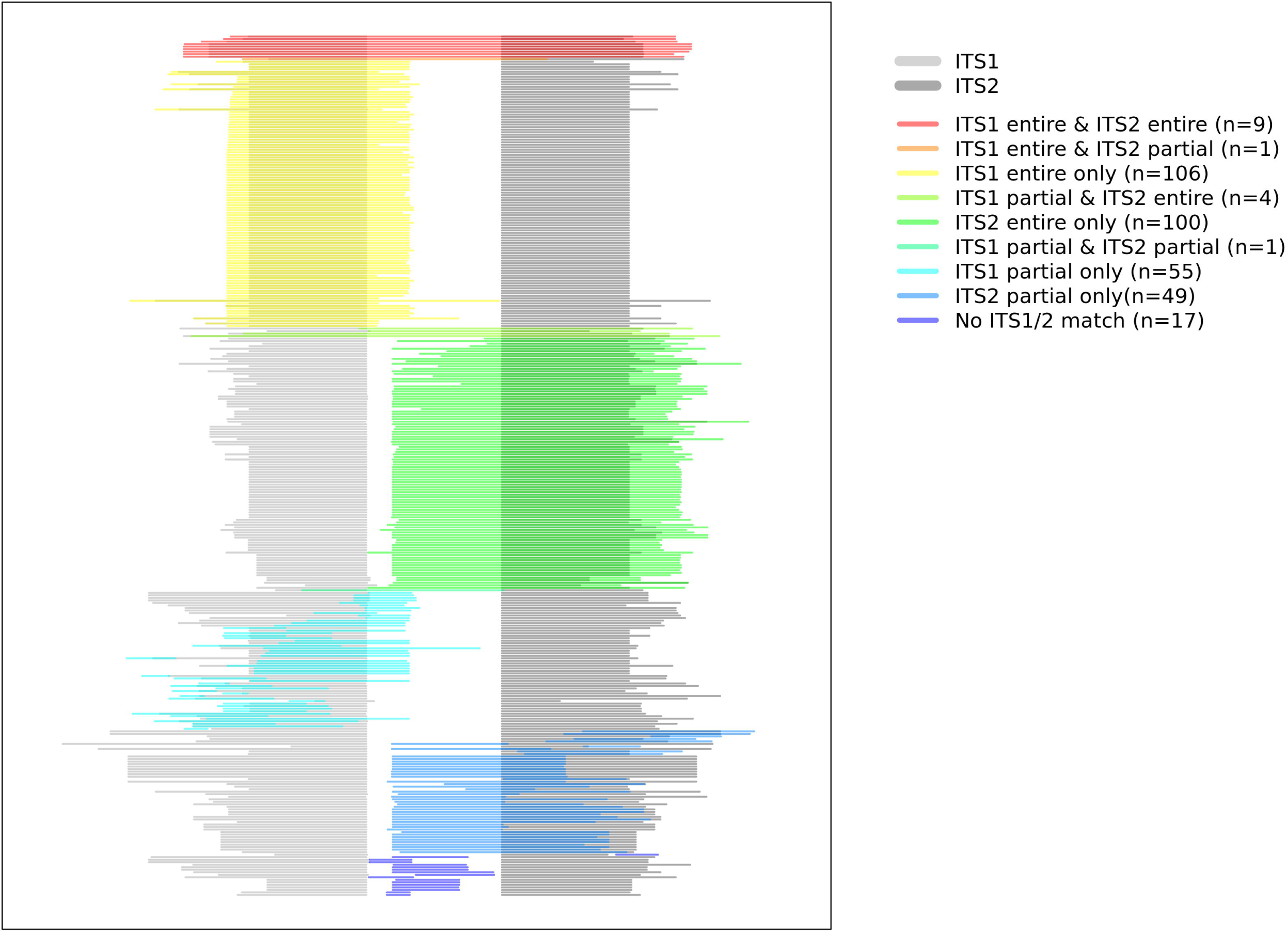
Summary of the number of DNA sequences identified per case.

**Figure 2.**
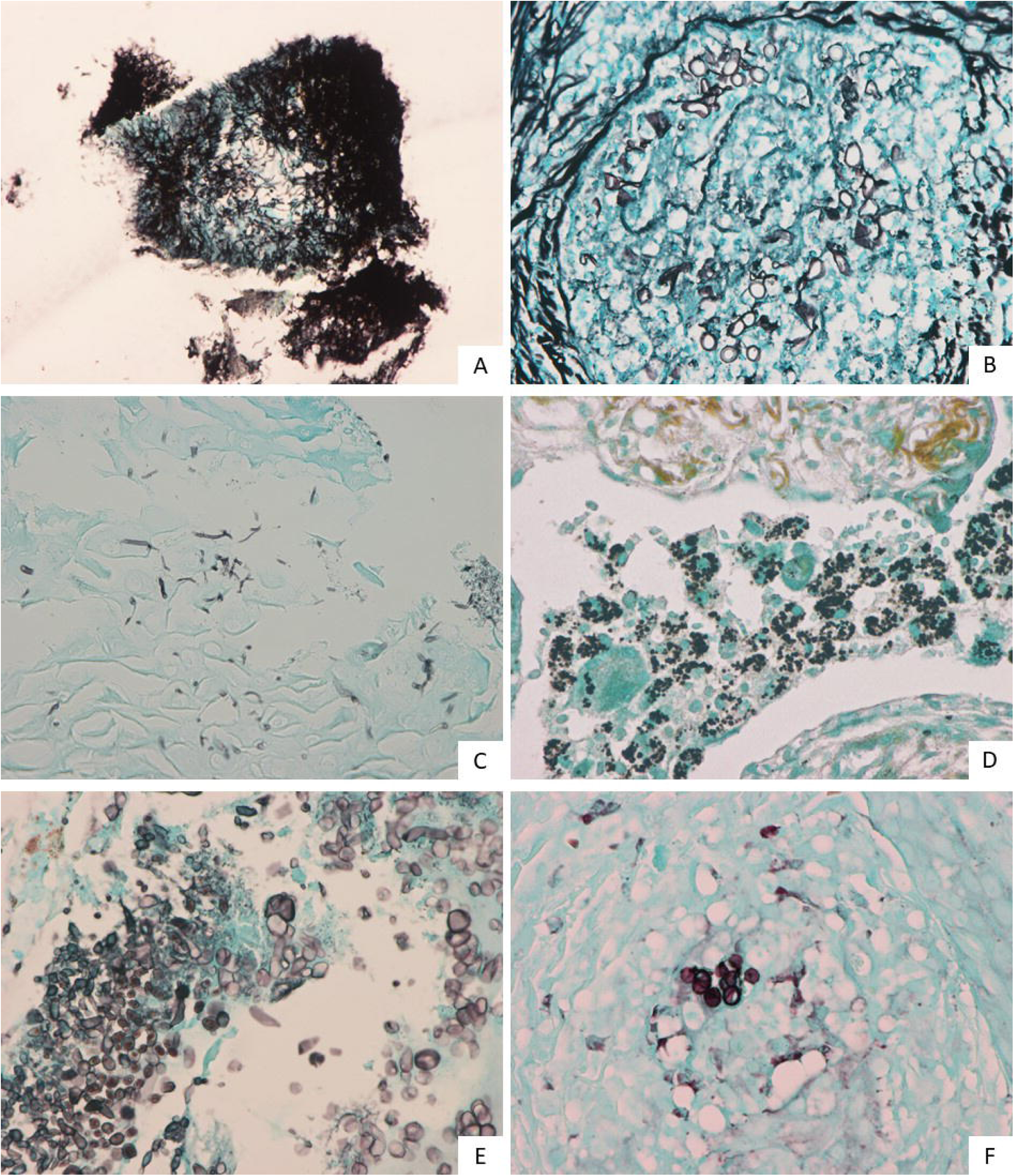
Visualization of nucleotide sequences used for fungal identification according to the degree of ITS1 and ITS2 sequence matching (the nucleotide sequences of the reference database were aligned with the starting base of the ITS2 region).

The mean storage period of fungal and no fungal cases was 4.2 years and 4.5 years, respectively (p = 0.643). The DNA concentrations measured by NanoDrop after DNA extraction in the groups with and without fungal DNA detection were 71.2 ng/ μ𝓁 and 95.5 ng/ μ𝓁, respectively, with no statistical significance (p=0.417). The fungal DNA detection rate was not statistically significant (p=0.376)

Representative microscopic images of some cases with inconsistent pathologic diagnosis and mycobiome analysis are summarized in Figure 3. *Aspergillus* and *Candida* species were detected in case 1-01 (Figure 3A). Compared to *Aspergillus* species commonly found in nasal cavities, thinner hyphae were observed, which may be misleading as sulfur granule of *Actinomyces. Aspergillus* and *Acremonium* species were detected in case 1-03 (Figure 3B). Contrary to case 1-01, it is thought to be misdiagnosed as *Mucor* species because of its slightly wider hyphae than *Aspergillus* species. Case 1-08 (Figure 3C) was also misdiagnosed as *Actinomyces*, which is thought to be similar to case 1-01. *Alternaria, Fusarium* and *Aspergillus* species were commonly detected. In case 2-08 (Figure 3D), yeast-form fungi were observed in alveolar macrophage on Gomori Methenamine-silver stain and *Starmerella cellae* was identified. *Starmerella cellae* is a relatively recently identified ovoid to ellipsoidal fungus.^19^ Case 2-09 (Figure 3E) is a fungus found in pharynx, which shows morphological findings different from those of *Candidda, Aspergillus*, and *Mucor*. In other words, yeast-form or short branching-type, fungal nuclei were found inside fungi like *Pneumocystis jiroveci. Cladosporium coloradense* was identified. Case 3-04 (Figure 3F) is a yeast-form fungus found in subcutaneous tissue and *Candida glabrata* were frequently identified by sequencing.

**Figure 3.**
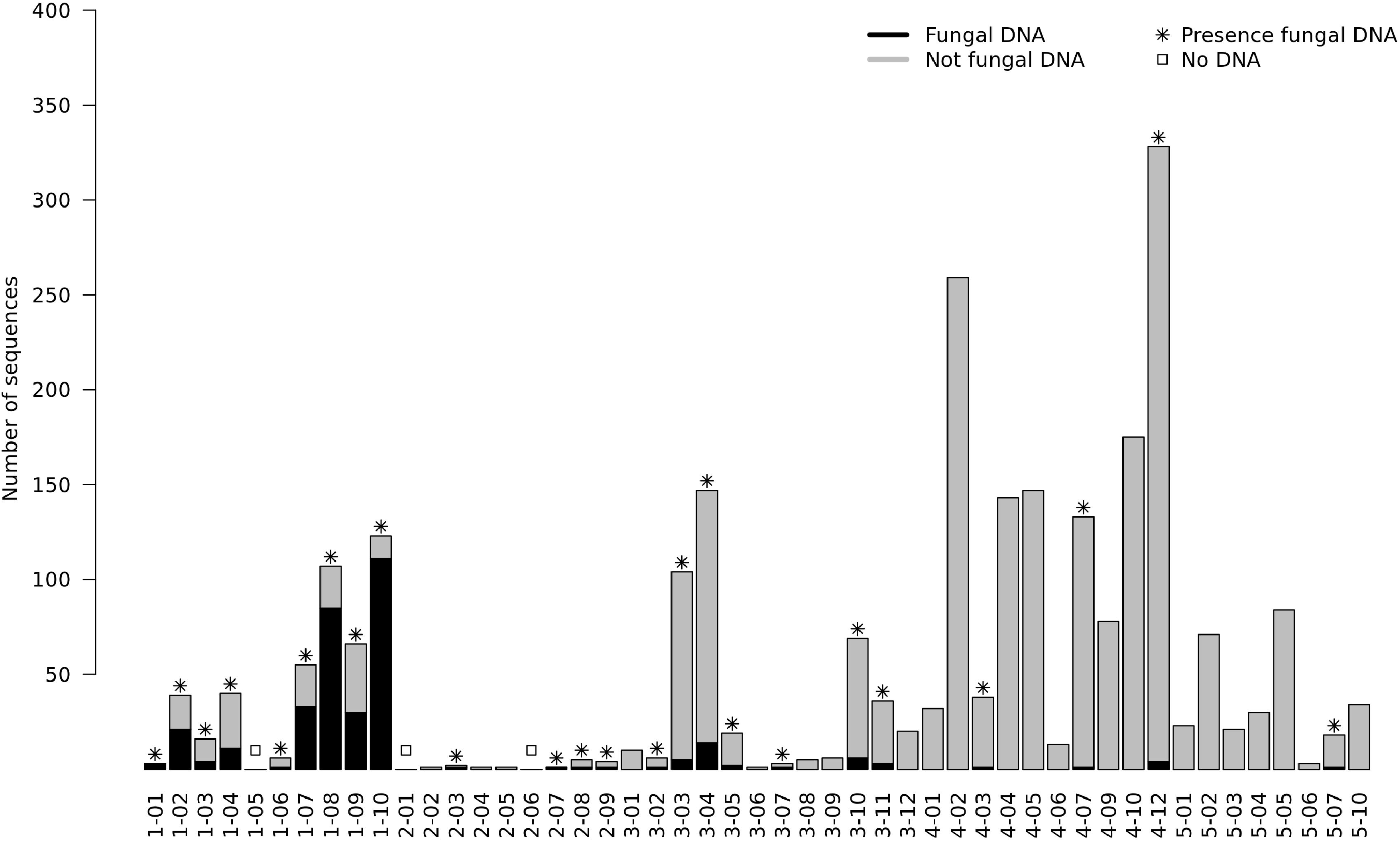
Representative microscopic images with inconsistent pathological and molecular diagnoses (magnification x600, case 1-01, 1-03, 1-08, 2-08, 2-09 and 3-04 as ordered by appearance).

All three cases diagnosed with *Actinomyces* were found to have fungi (cases 1-01, 1-02 and 1-08). Case 3-02 was diagnosed as *Blastomyces dermatitidis* by sequencing analysis (summary of sequencing results: 575 nucleotide sequences of U18364.1 were identical without gap opening, and 549 nucleotide sequences of EF592163.1 were identical without gap opening, respectively). One strand of fungal DNA was identified from the DNA extracted from the FFPE tissue, and the BLASTN analysis showed the highest probability match was *Candida africana* with a bit score of 468. In this case, the bit score was 154 based on the separate sequence of *Blastomyces dermatitidis*. Among three cases of pathologically diagnosed cryptococcosis, one case (case 3-10) was found to have a nucleotide sequence of *Cryptococcus neoformans* with a bit score of 296. In case 4-12, the nucleotide sequence of *Candida africana* with a bit score of 438 was suggested with the possibility of *Pneumocystis jiroveci* with a bit score of 73.1. Four cases of pathologically unidentifiable fungi (cases 2-03, 2-08, 2-09 and 3-04) were found to most likely be *Acremonium acutatum, Starmerella cellae, Cladosporium coloradense*, and *Candida glabrata*, respectively.

## Discussion

We performed mycobiome analysis in fungal infected FFPE tissues using nanopore sequencing. The detected fungal DNA occupied approximately one-third of the ITS1 entire region, the ITS2 entire region, and other regions, respectively. The advantages of nanopore sequencing compared with Sanger sequencing are as follows. First, unlike Sanger’s method, which requires a lot of DNA, nanopore sequencing can be performed with a small amount of DNA (in theory, even with one strand). Second, nanopore sequencing method could sequenced DNA separately, even if the sample contains a variety length of DNA.^20^ Third, nanopore sequencing equipment (i.e., MinION) could be operated at lower cost (about $1,000) compared to Sanger sequencing equipment. This low initial cost is a critical factor in the introduction of equipment in small pathology laboratories. Compared to the 2nd generation sequencing equipment, nanopore sequencing has the advantage of sequencing with damaged DNA because there is no PCR amplification process in the sequencing process itself. In this study, about one third of the fungal DNA was not an ITS1 entire match or an ITS2 entire match. Therefore, these fungi can be effectively detected using nanopore sequencing.

It is necessary to find pathogenic fungi in various fungi detected by mycobiome analysis. It may be assumed that dominant fungi may have been associated with the disease at the site of fungal infection. In cases 1-07 and 1-10 diagnosed as esophageal candidiasis, *Candida* species comprise 72% and 93% of the fungal DNA. In such cases, it would be clear to diagnose that the infection is caused by *Candida* species. In case 1-09, however, only 7% of the fungal DNA was *Candida* species. Similarly, *Rhizopus* species were detected in *Mucor* infections, but only a small fraction of the fungal DNA. Considering these cases, it is considered desirable to include the process of detecting normal flora in adjacent uninfected sites and selecting pathogenic fungi associated with infection.

We performed multiplex PCR to increase fungal DNA concentration. However, 2,184 non-fungal DNAs accounted for 86.46% of the total DNA. Of the 2,184 DNAs not identified as fungi, only 382 (17.50%) were identified in human genome (GRCh38). Because ribosomal DNA is present in all living organisms, including fungi, humans, and bacteria, it may be difficult to amplify only the fungal ITS region. Nevertheless, we were only able to detect fungal DNA in 49.0% of cases. This detection rate is not high enough to be applied in actual clinical situation and needs improvement. We suggest the following about the causes of low fungal DNA detection rates. The first and most important is a low sequencing output. The third generation sequencing technique is known to have less sequencing output than the second generation sequencing equipment. Therefore, increasing the purity of the DNA to be sequenced is important for research. Since we used multiplex sequencing, we were forced to reduce the amount of DNA we analyzed per sample. We thought that the lack of sufficient DNA sequencing in each case was the main cause of the failure to detect fungal DNA. The second is the lack of fungal DNA evaluation methods. Unlike human cells, in which cell viability (closely related to DNA quality) could be predicted by evaluation of H&E stain slide, it is not known to evaluate the viability of fungi in H&E stain. Many studies have shown that inflammation and hypoxia are closely related, and hypoxia induced by inflammation caused by fungal infections may result in damage to fungal DNA and consequently affect the sequencing.^21^ Third, the aforementioned ribosomal DNA is present in almost all living things. We thought this could be overcome by using microdissection, which extracts DNA from fungi as much as possible. Fourth, there is a lack of high quality DB. Even in the database of NCBI RefSeq Targeted Loci Project, there is only one species (*murina*) in the *Pneumocystis* genus. In other words, to diagnose *Pneumocystis jiroveci*, a separate DB should be constructed. In addition, when we analyzed the GenBank file, there were 5,602 ITS1 full sequences and 7,552 ITS2 full sequences in 11,133 genus. Fifth, it is known that 50 μm thickness of tumor biopsy specimens can sufficiently perform NGS for diagnostic purposes.^22^ Our study was also performed with a 50 μm thickness, referring to the results of this study. However, if more tissues (e.g., 100 μm thick) were used, the fungal detection rate could be increased.

*Candida albicans* is the most commonly isolated *Candida* species in the clinical setting. Of the 162 *Candida* species, 141 *Candida africana* (87.04%) and 8 *Candida albicans* (4.94%) were detected. These two *Candida* species are similar in ITS sequences. Comparing the ITS sequences (ITS1+5.8S+ITS2) of *Candida africana* (NR_138276.1, 447bp) based on *Candida albicans* (NR_125332.1, 446bp) shows 99.192% identity. There is a 1 bp gap open at ITS1 in *Candida albicans*. Two mismatched nucleotides are located in ITS2, where the DNA base “C” in *Candida albicans* is “T” in *Candida africana*. It is generally known that nanopore sequencing has a slightly higher error rate than second-generation NGS sequencing.^23^ DNA extracted from FFPE tissue is known to have a higher C:T conversion than DNA extracted from fresh frozen tissue.^24^ Because of these, our method seems to be difficult to identify precise fungi, especially at the species level.

In summary, we have identified the possibility of finding pathogenic fungi through mycobiome analysis in fungal infected FFPE tissues. However, we have found problems to be solved for further studies, such as increasing sequencing output, increasing fungal DNA concentration, excluding normal flora, and expanding fungal DB. If we develop a method to characterize pathogenic fungi in FFPE tissues in a follow-up study, we think it will help patients to use appropriate antifungal agents.

## Author Contributions

Conceptualization: SJB

Data curation: SJB, TBL, HYN, JYC, SHP

Formal analysis: SJB

Funding acquisition: SJB, KCC

Investigation: SJB

Methodology: SJB

Project administration: SJB

Resources: KMK, HSL, SHP, JYC

Software: SJB

Supervision: SJB

Validation: SJB

Visualization: SJB

Writing - original draft: TBL, HYN

Writing - review & editing: SJB

## Supplementary legends

Supplementary Table 1. The clinicopathological information of remaining cases which fungal DNA was not sequenced.

Supplementary Table 2. The result of BLAST program.

Supplementary Figure 1. All available representative of pathologic images in this study.

