## Supplementary Table 1 for "Mycobiome analysis in fungal Infected formalin-fixed and paraffin-embedded tissues for identification of pathogenic fungi: A pilot study"

Supplementary Table 1. The basic clinicopathological information of no-fungal DNA detected cases (sorted by pathologic diagnosis).

| Case | Sex/Age | Location | Pathology diagnosis | Number of sequences |
| --- | --- | --- | --- | --- |
| 2-01 | M/51 | Stomach | Actinomyces | 0 (No DNA) |
| 2-02 | F/40 | Eye | Actinomyces | 1 |
| 2-05 | F/42 | Paranasal sinus | Aspergillosis | 1 |
| 2-06 | F/58 | Paranasal sinus | Aspergillosis | 0 (No DNA) |
| 3-01 | M/60 | Lung | Aspergillosis vs. Mucormycosis | 10 |
| 4-10 | M/73 | Paranasal sinus | Aspergillosis vs. Mucormycosis | 175 |
| 5-03 | M/88 | Paranasal sinus | Aspergillosis vs. Mucormycosis | 21 |
| 5-04 | M/64 | Lung | Aspergillosis vs. Mucormycosis | 30 |
| 2-04 | M/52 | Stomach | Candida | 1 |
| 3-08 | M/63 | Lung | Cryptococcus | 5 |
| 3-09 | F/51 | Lung | Cryptococcus | 6 |
| 5-05 | M/61 | Lung | Cryptococcus | 84 |
| 5-06 | F/60 | Lung | Cryptococcus | 3 |
| 5-10 | F/11 | Liver | Cryptococcus | 34 |
| 5-01 | M/58 | Terminal ileum | Histoplasmosis | 23 |
| 5-02 | M/58 | Adrenal gland | Histoplasmosis | 71 |
| 1-05 | F/79 | Paranasal sinus | Mucormycosis | 0 (No DNA) |
| 3-06 | M/70 | Orbit | Mucormycosis | 1 |
| 3-12 | F/14 | Lung | Mucormycosis | 20 |
| 4-01 | M/44 | Lung | Mucormycosis | 32 |
| 4-02 | M/56 | Lung | Mucormycosis | 259 |
| 4-04 | F/50 | Lung | Mucormycosis | 143 |
| 4-05 | F/4 | Lung | Pneumocystis | 147 |
| 4-06 | M/5 | Lung | Pneumocystis | 13 |
| 4-09 | M/48 | Lung | Pneumocystis | 78 |
