## Supplementary figures and images for "Mycobiome analysis in fungal Infected formalin-fixed and paraffin-embedded tissues for identification of pathogenic fungi: A pilot study"

### Supplementary Figure 1

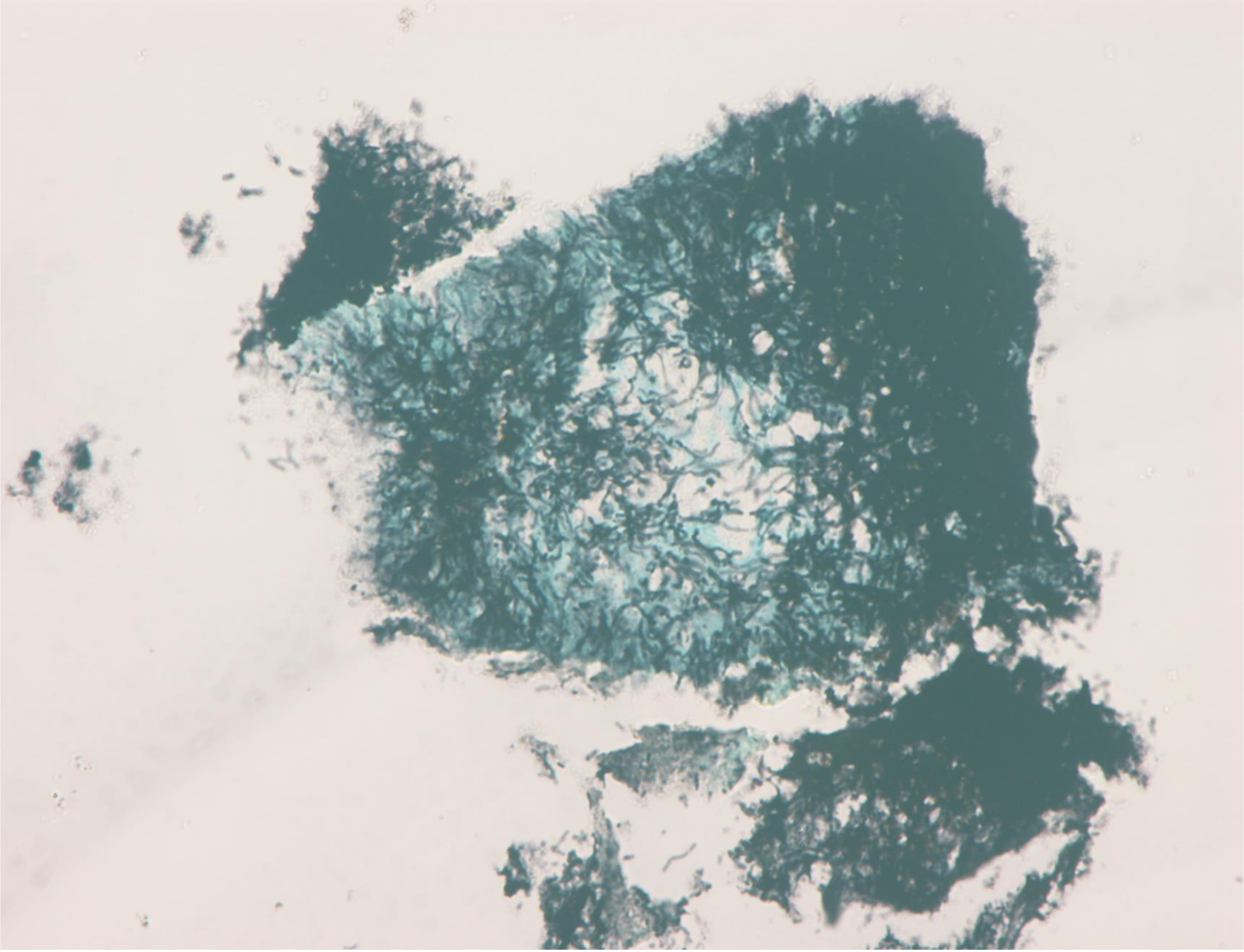

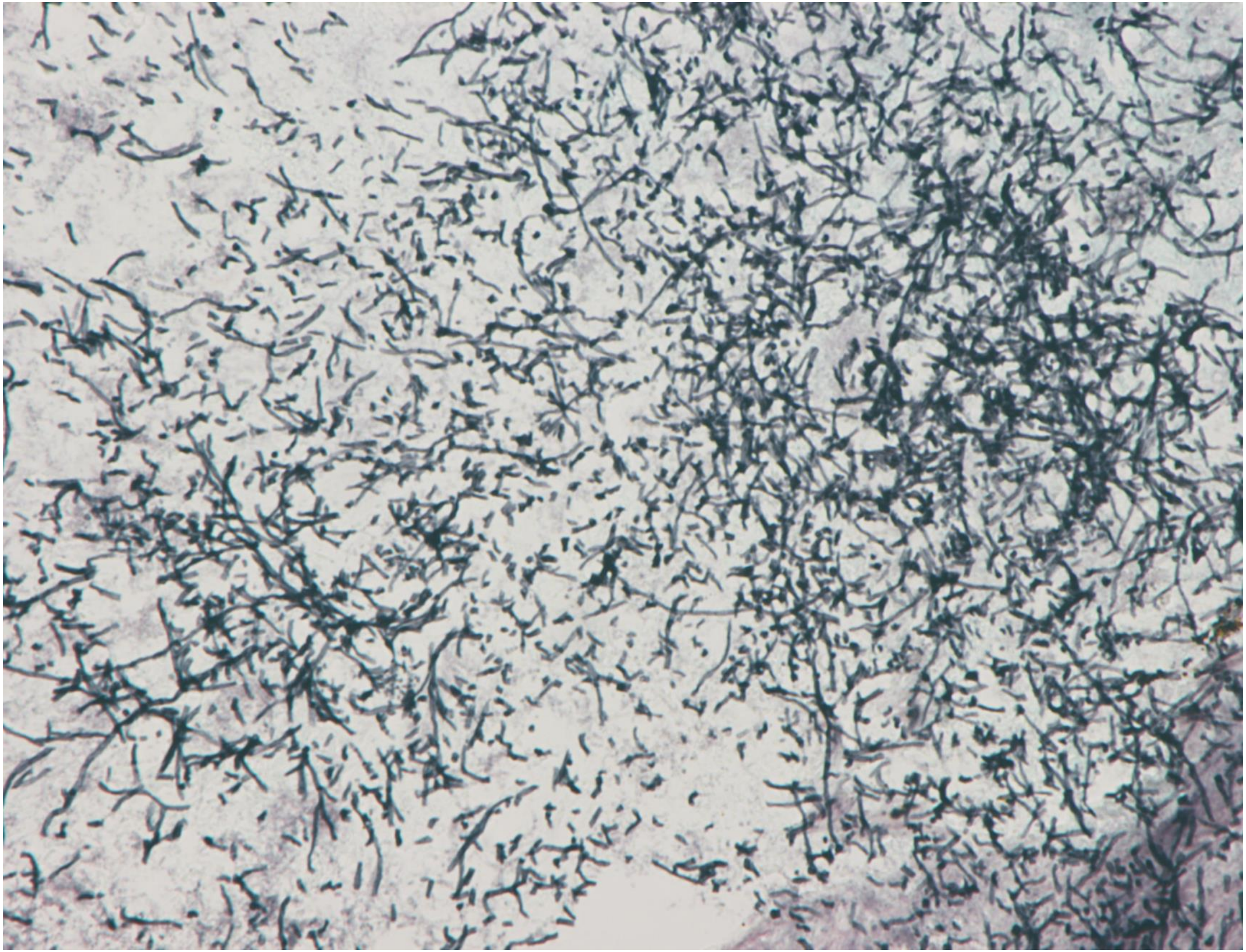

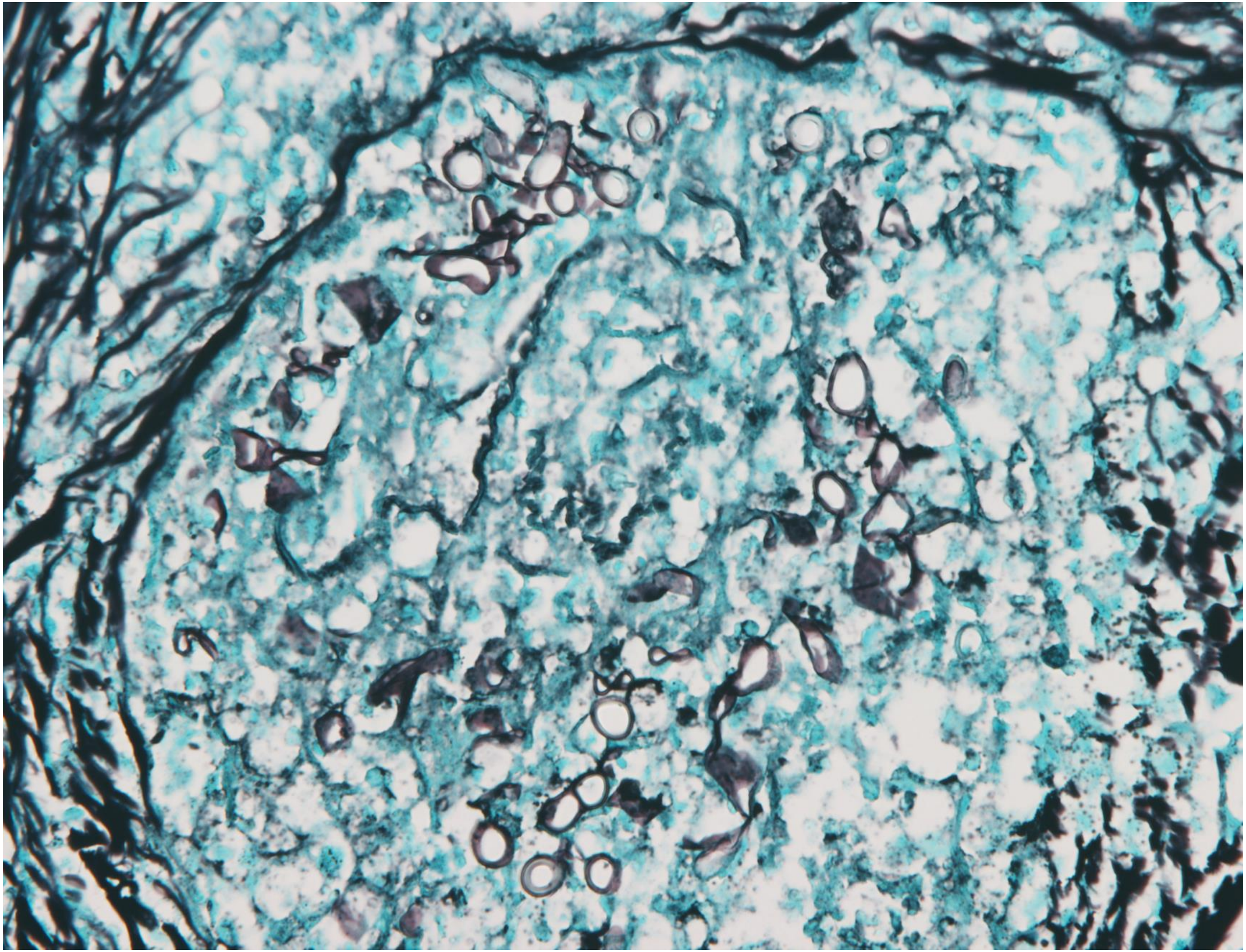

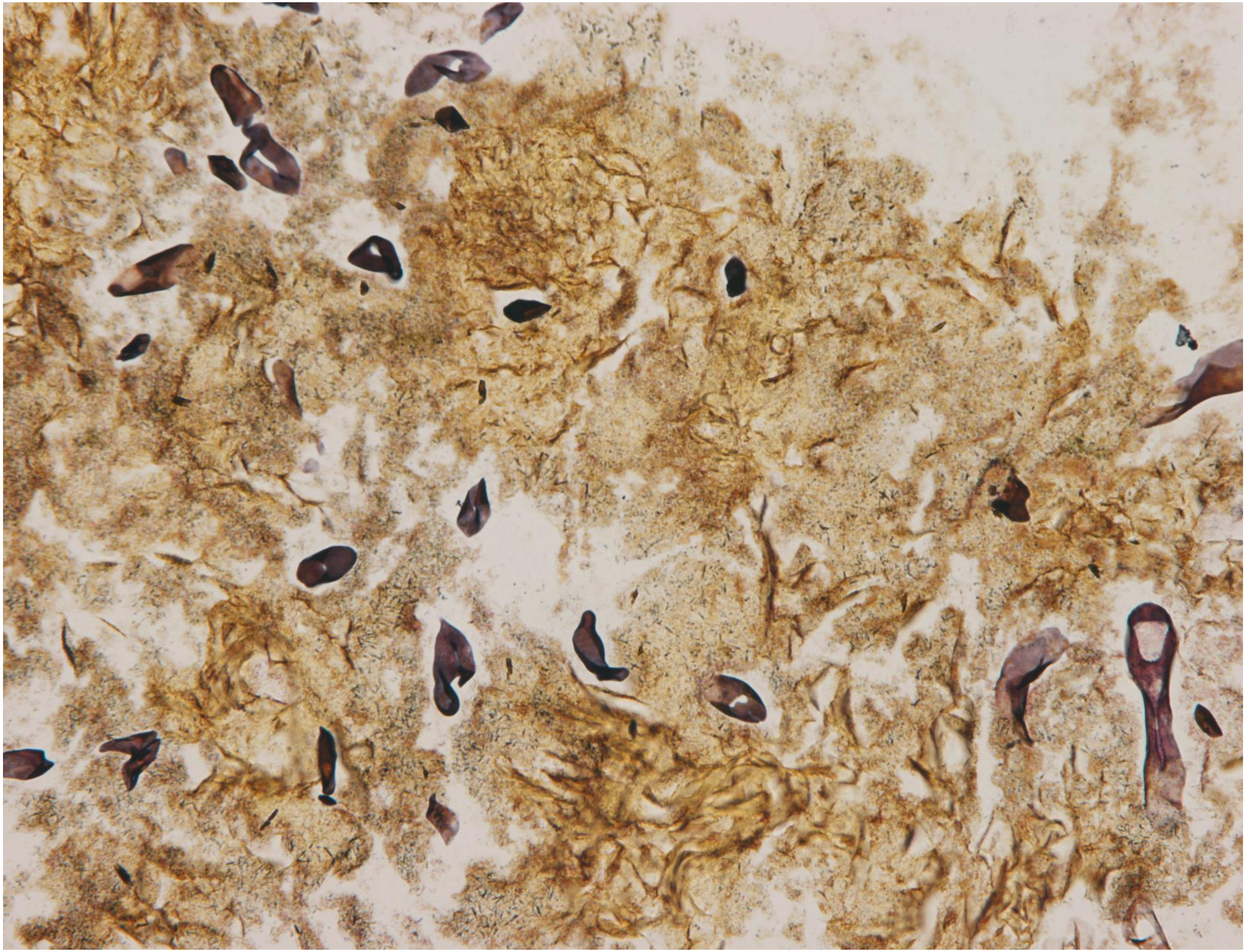

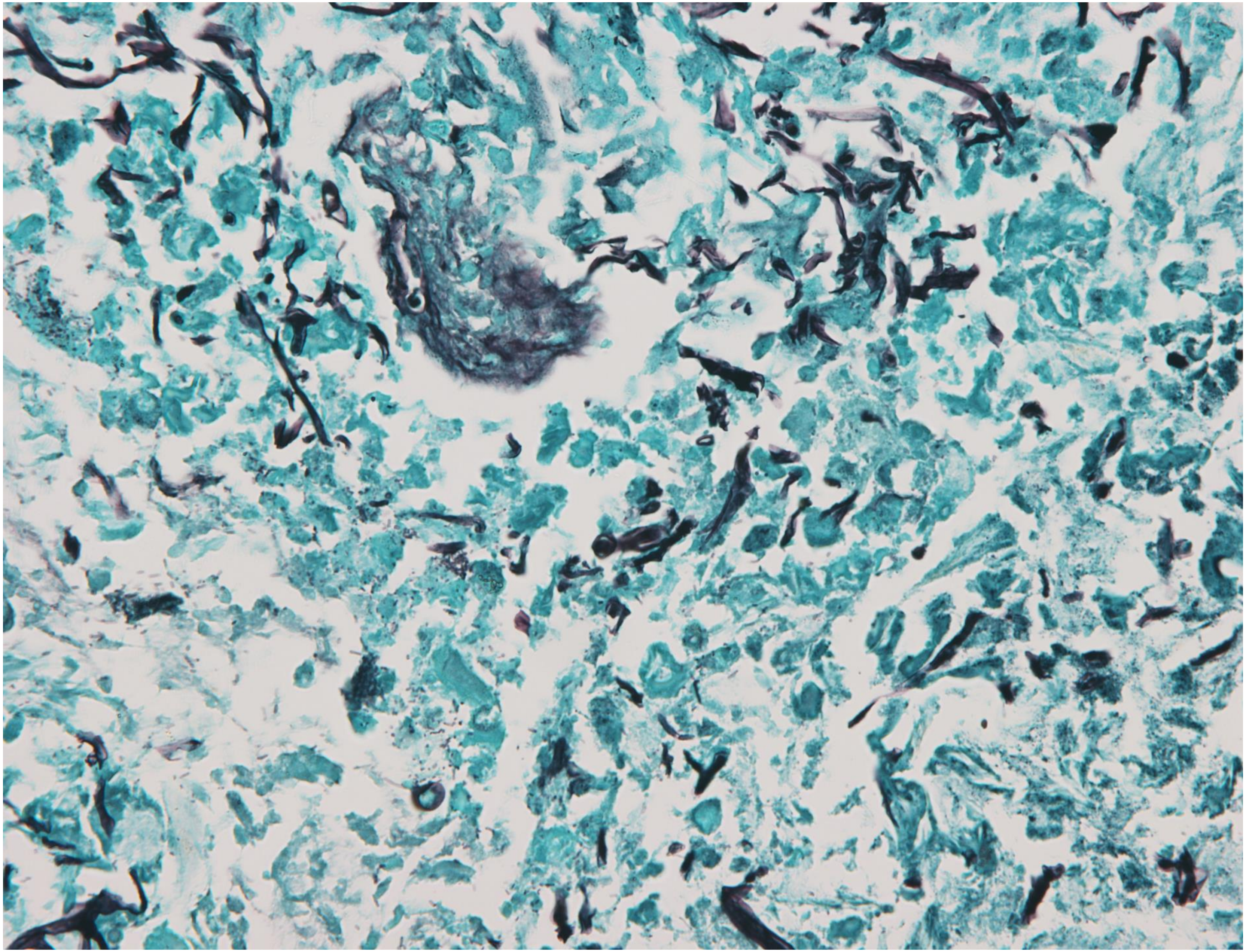

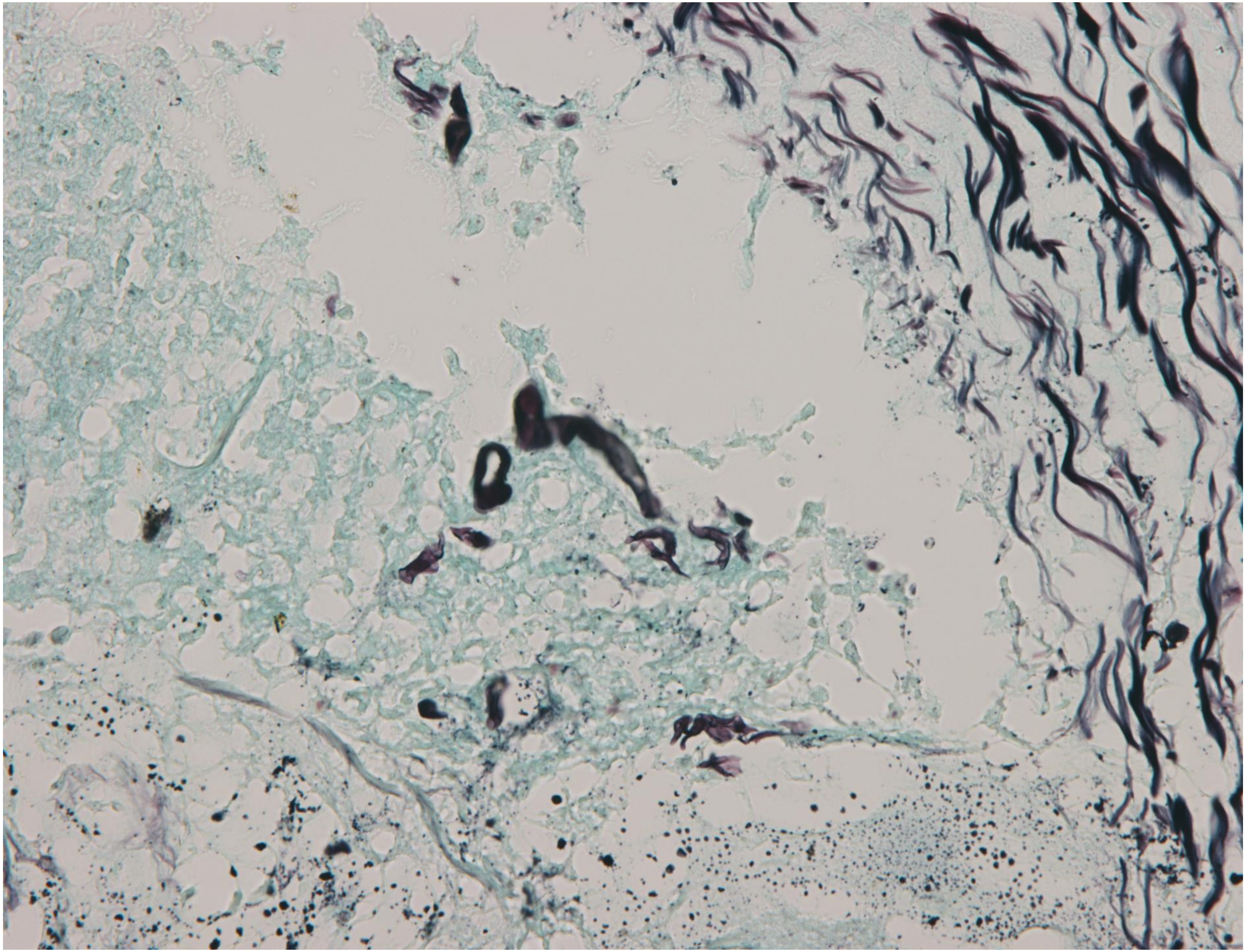

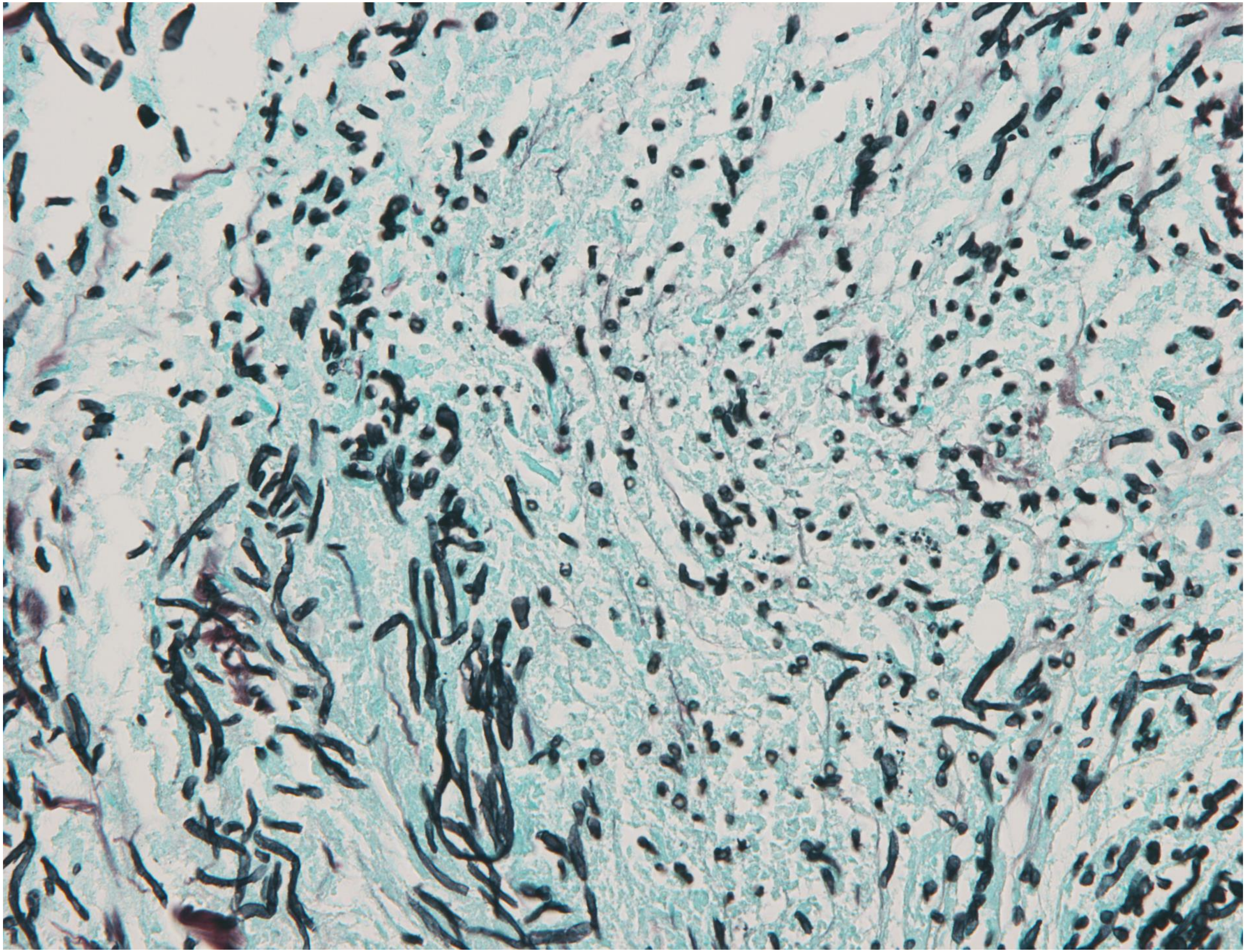

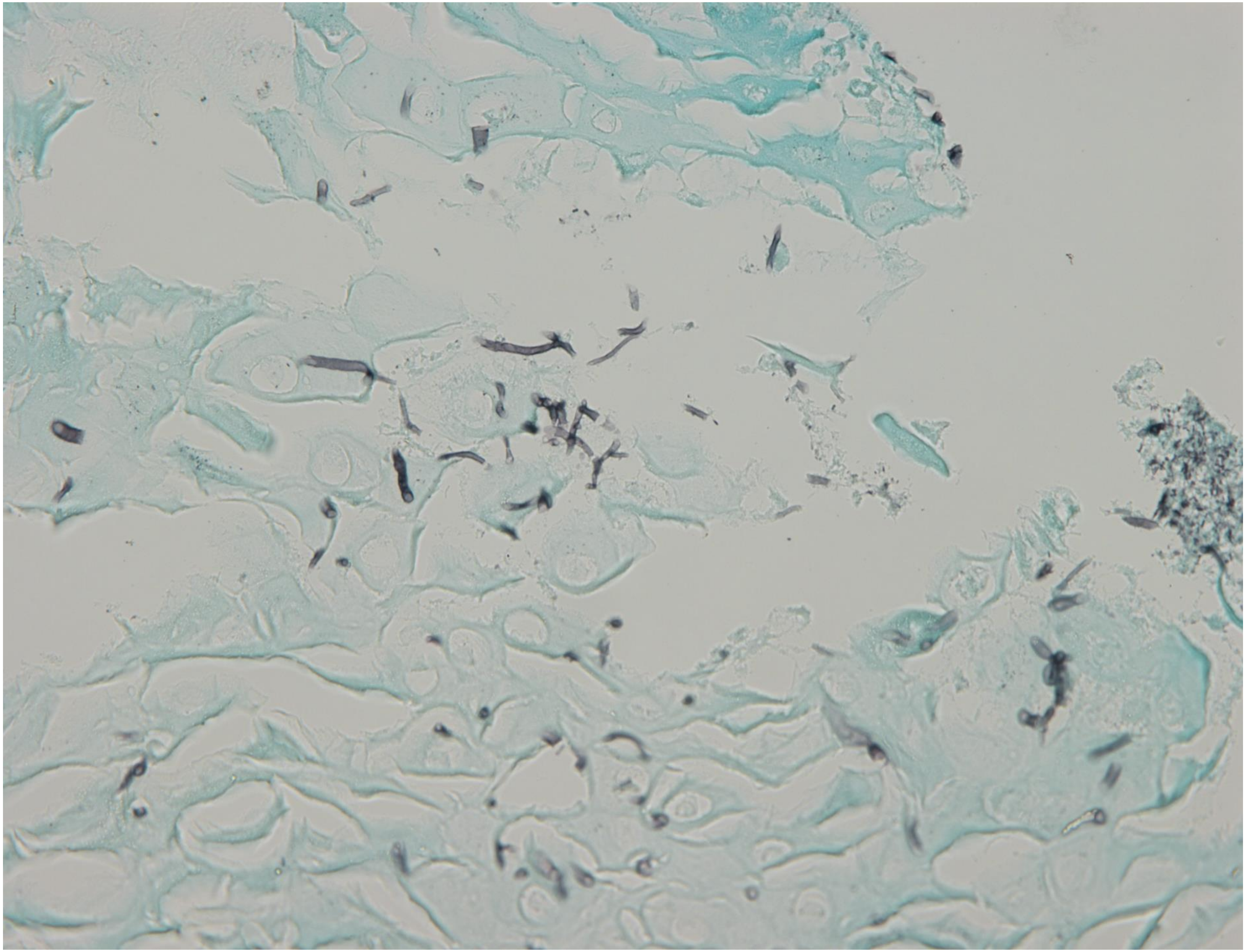

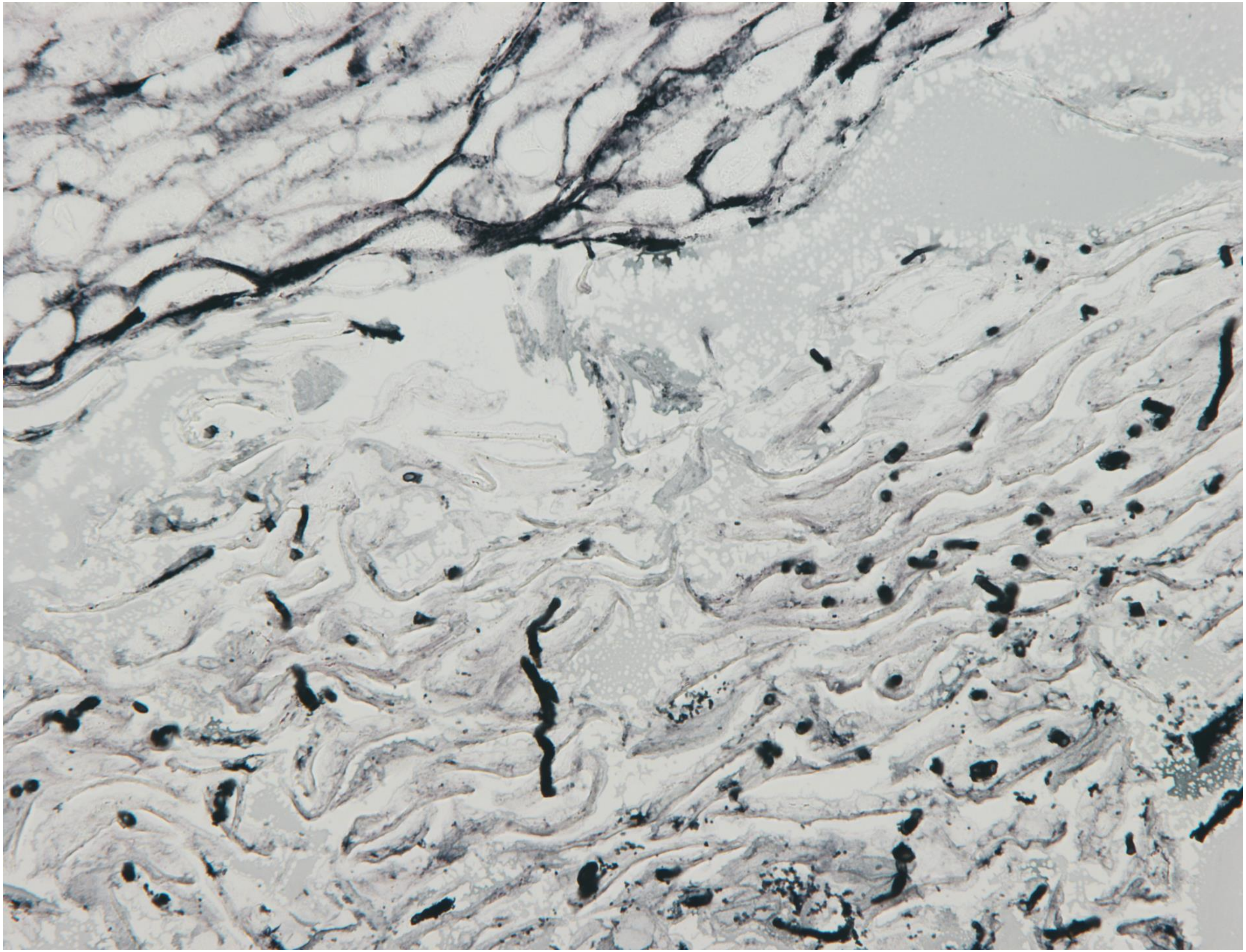

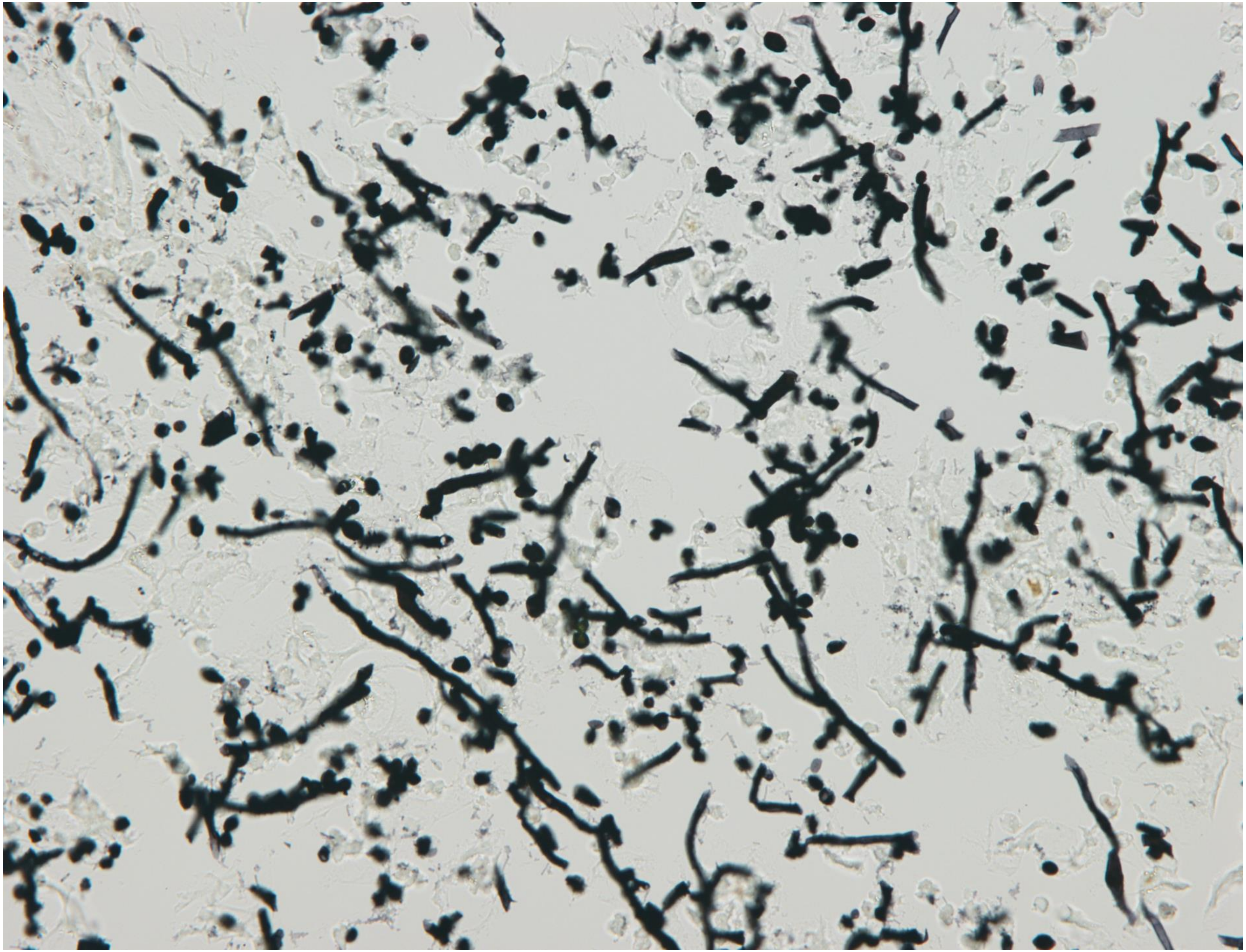

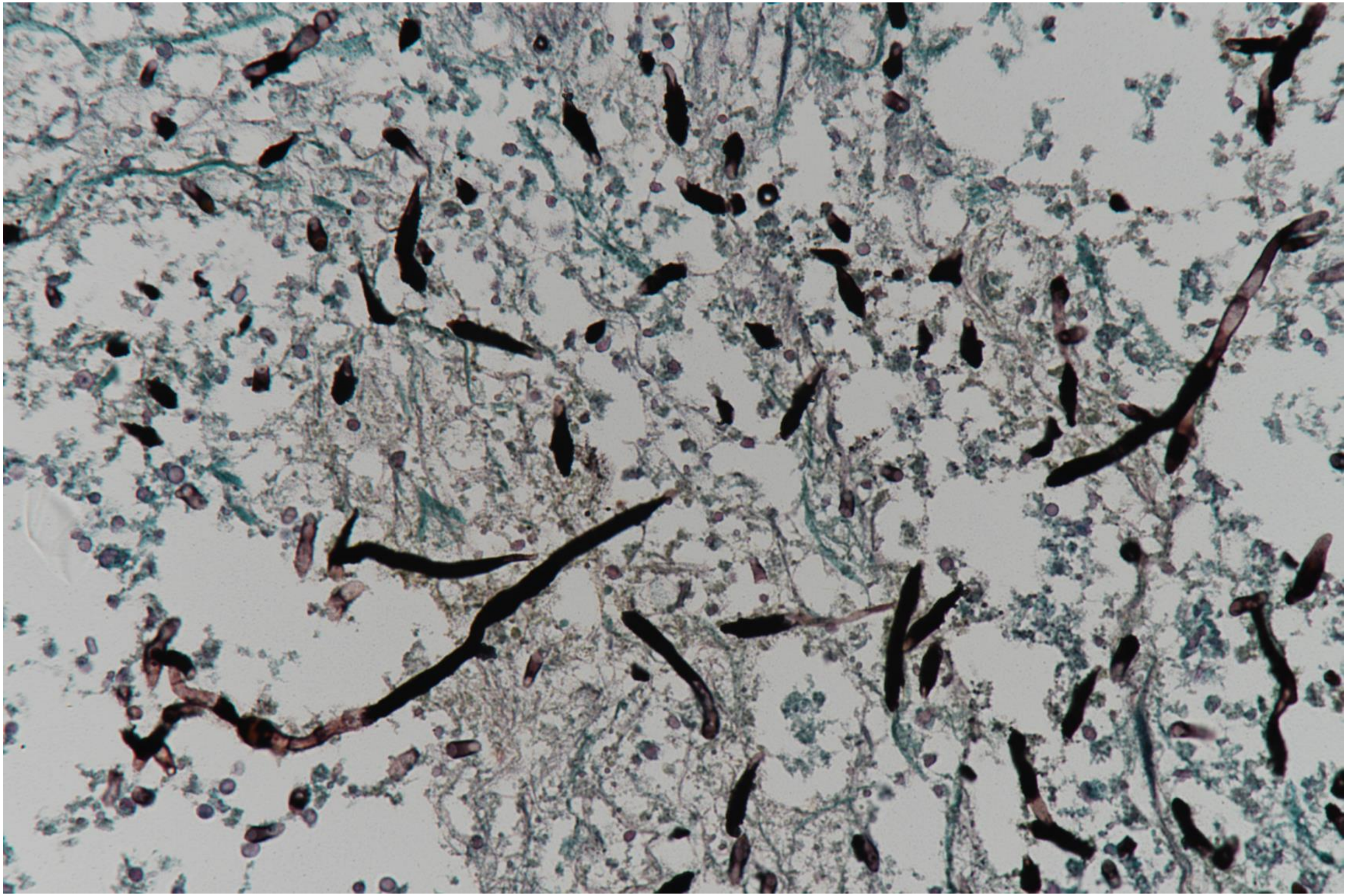

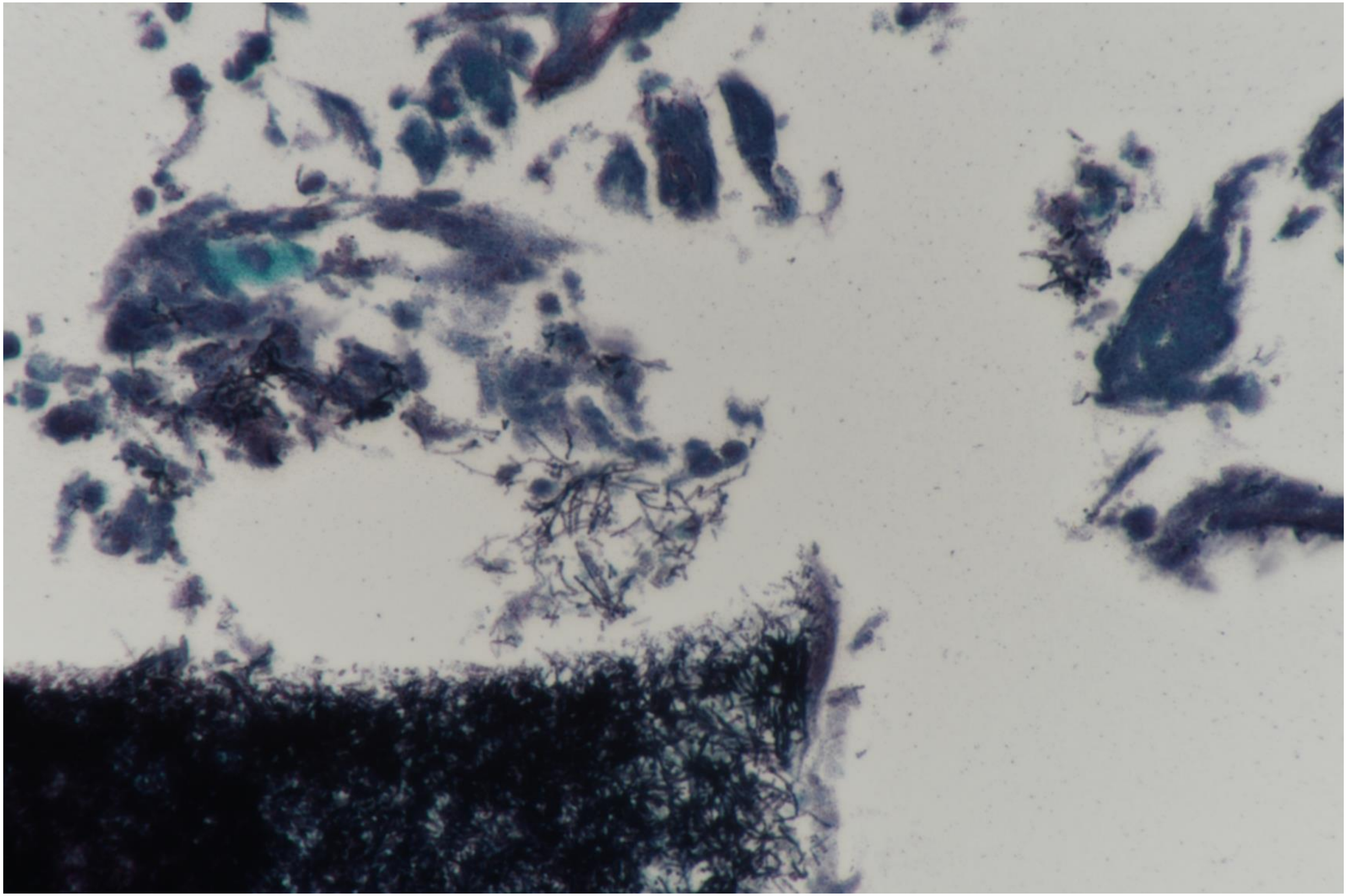

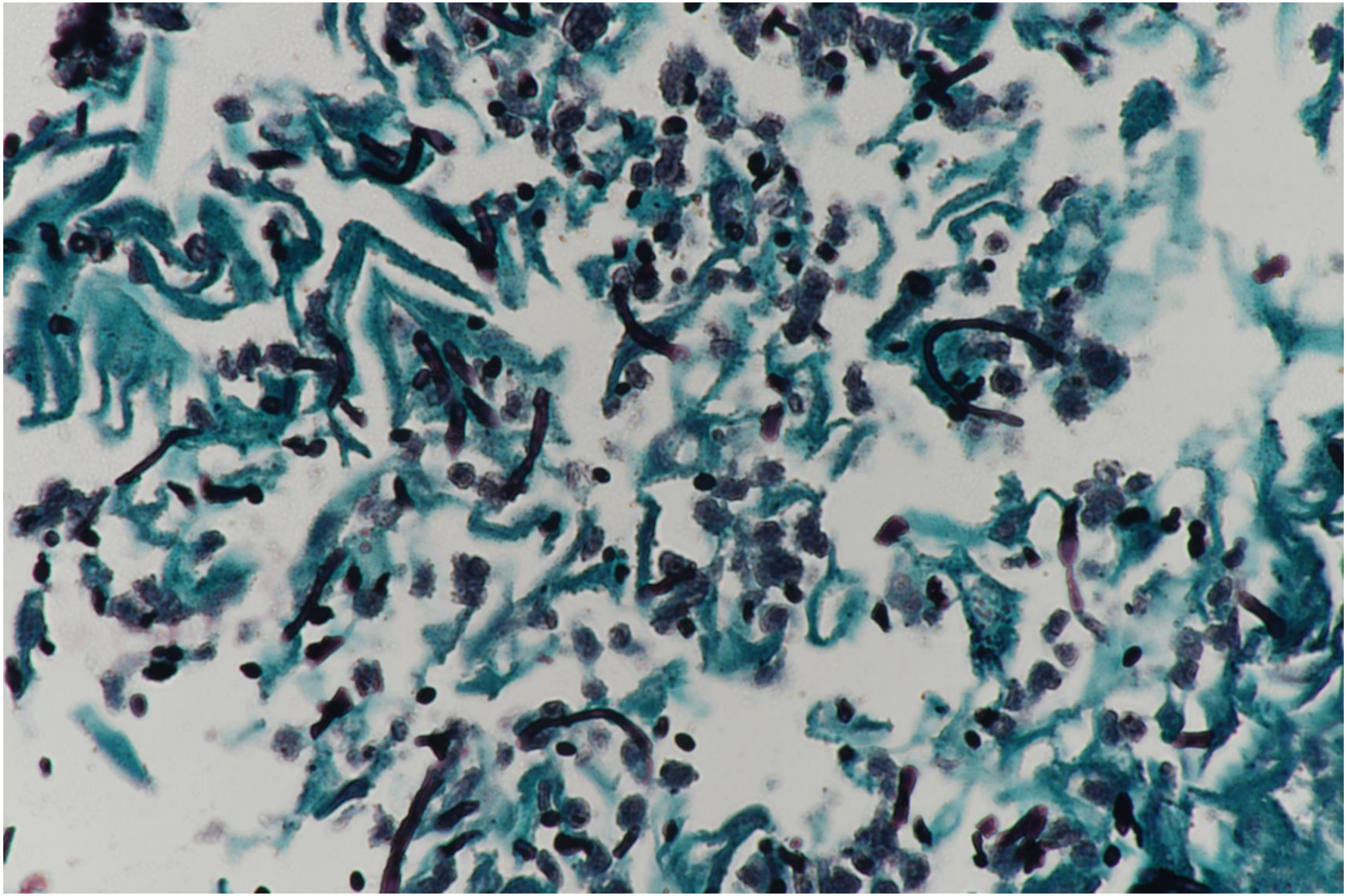

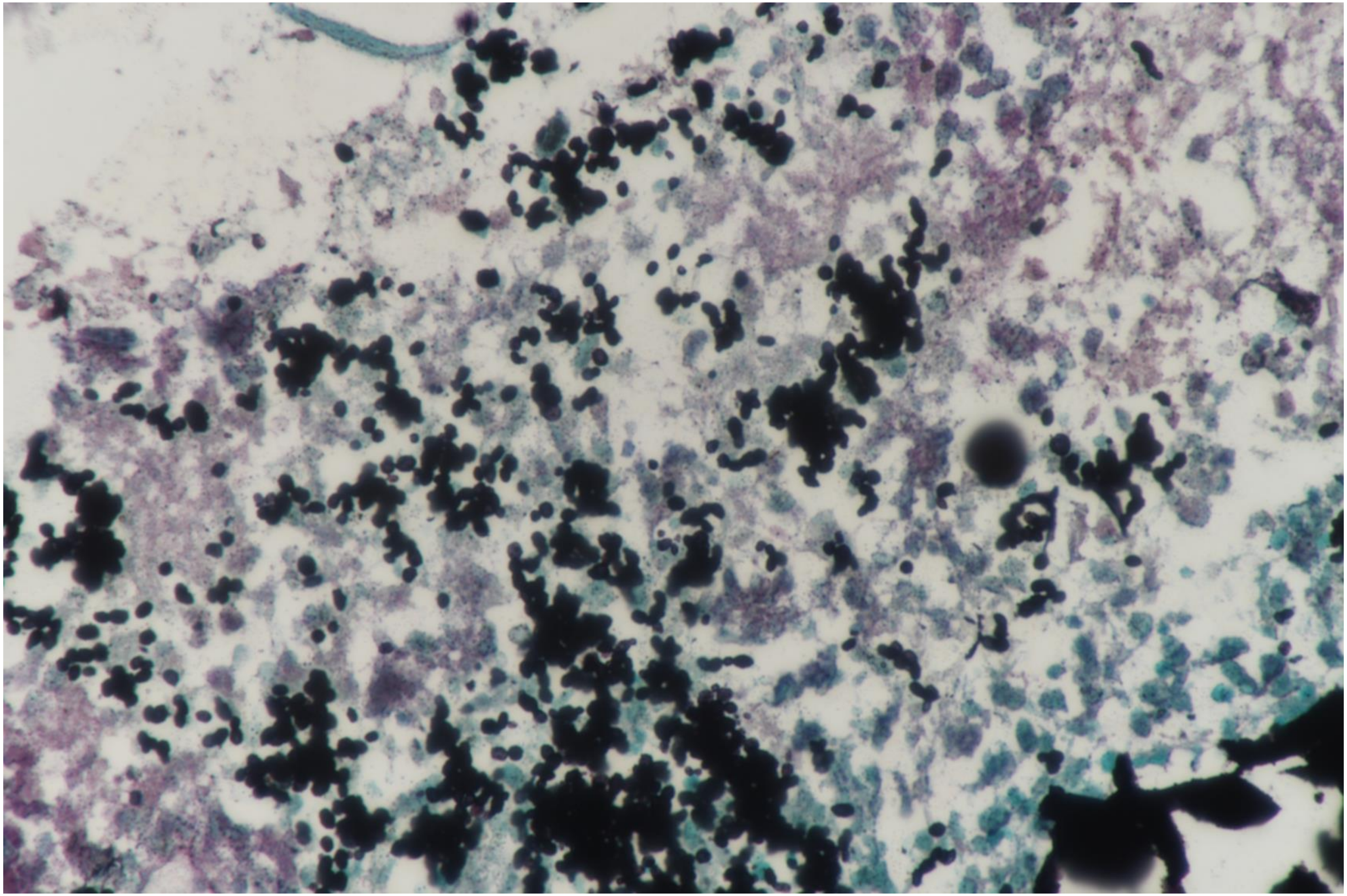

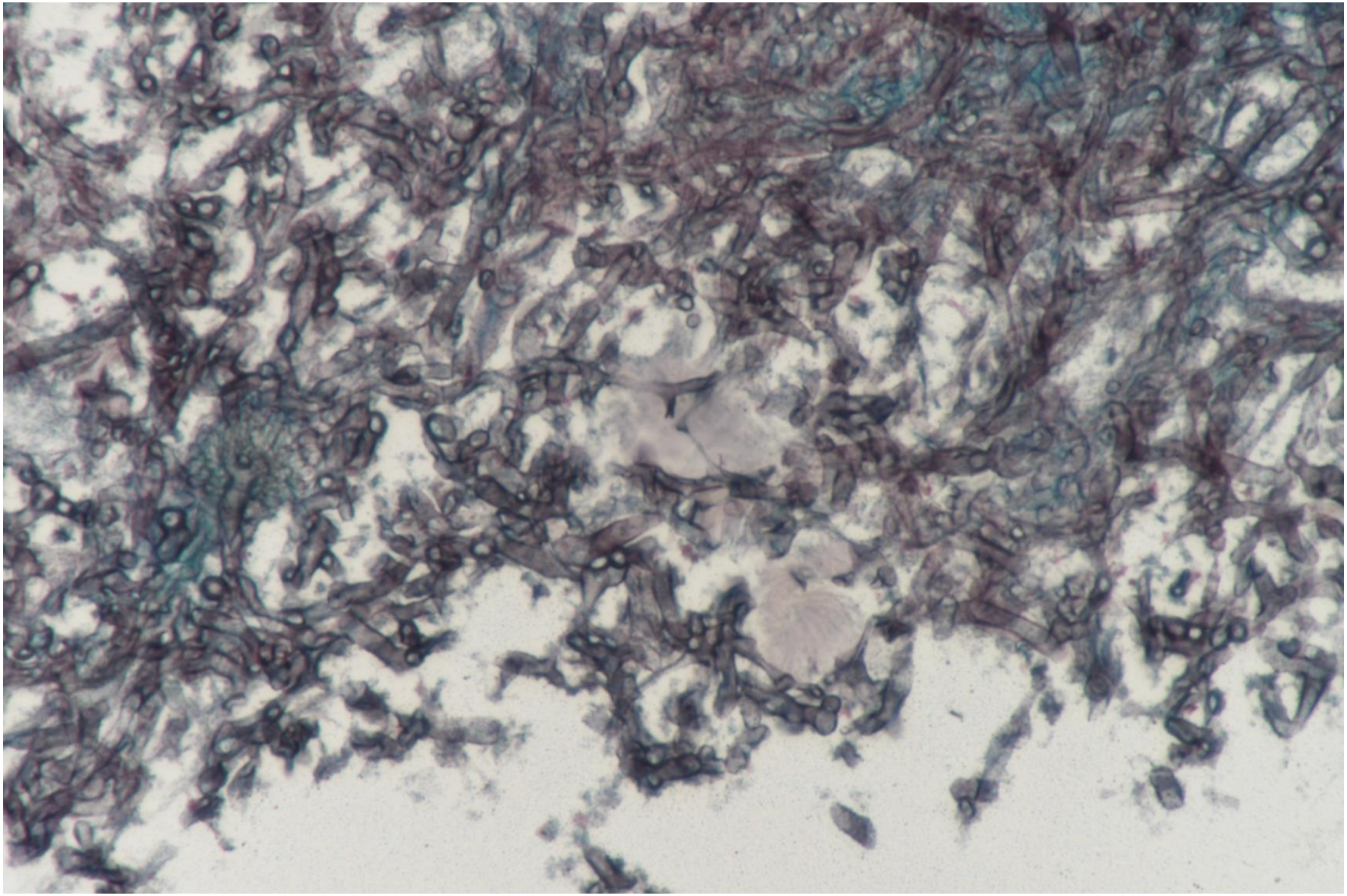

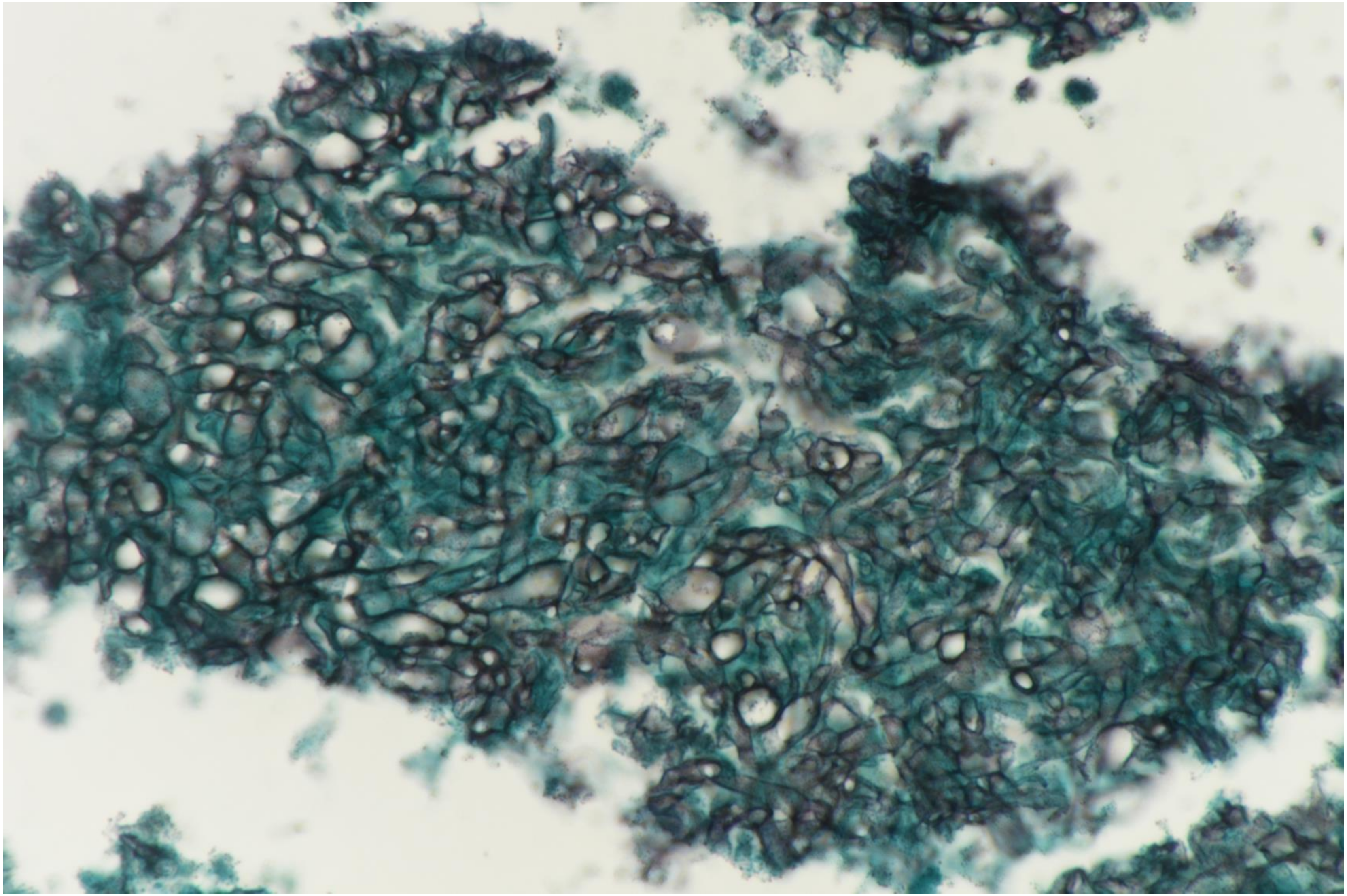

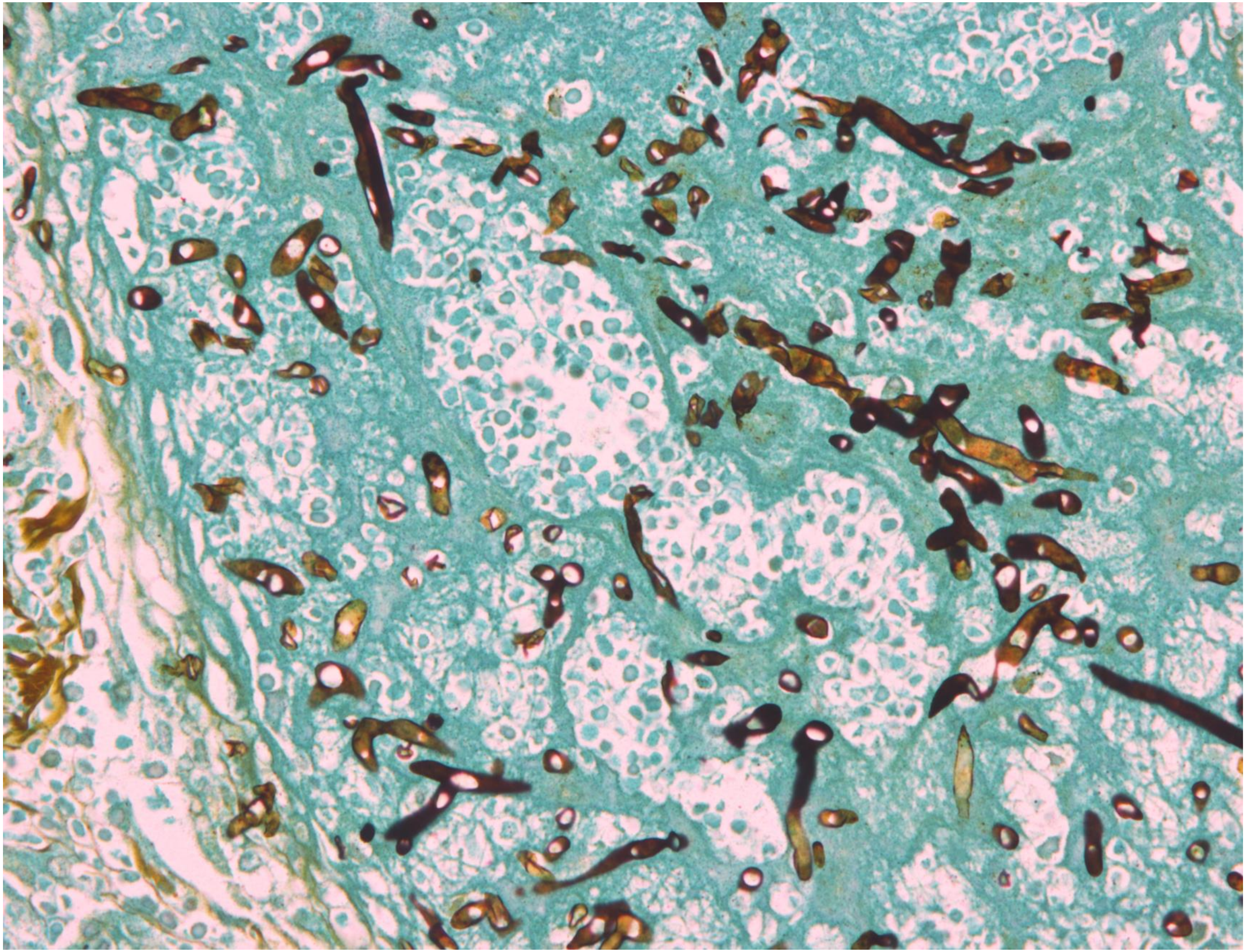

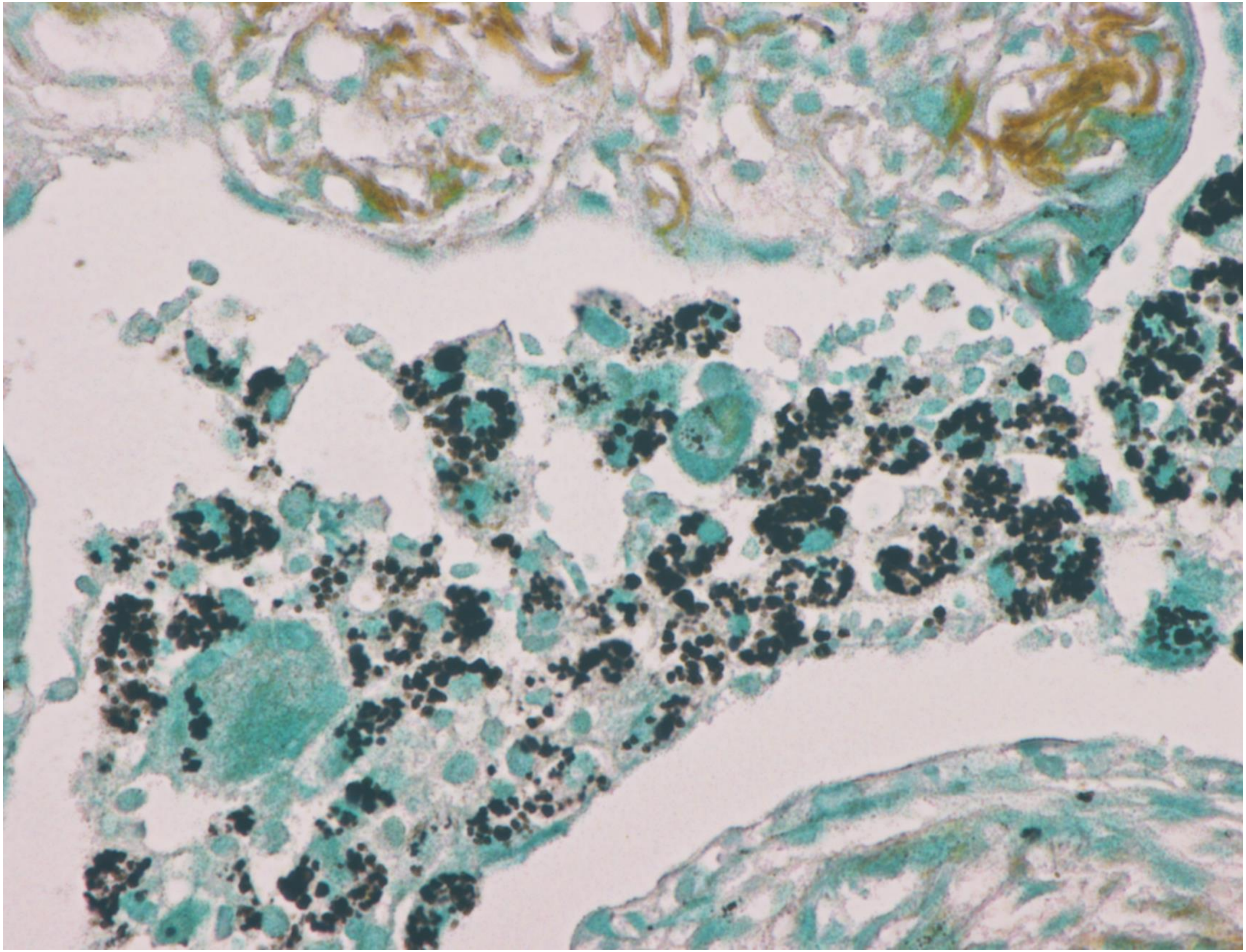

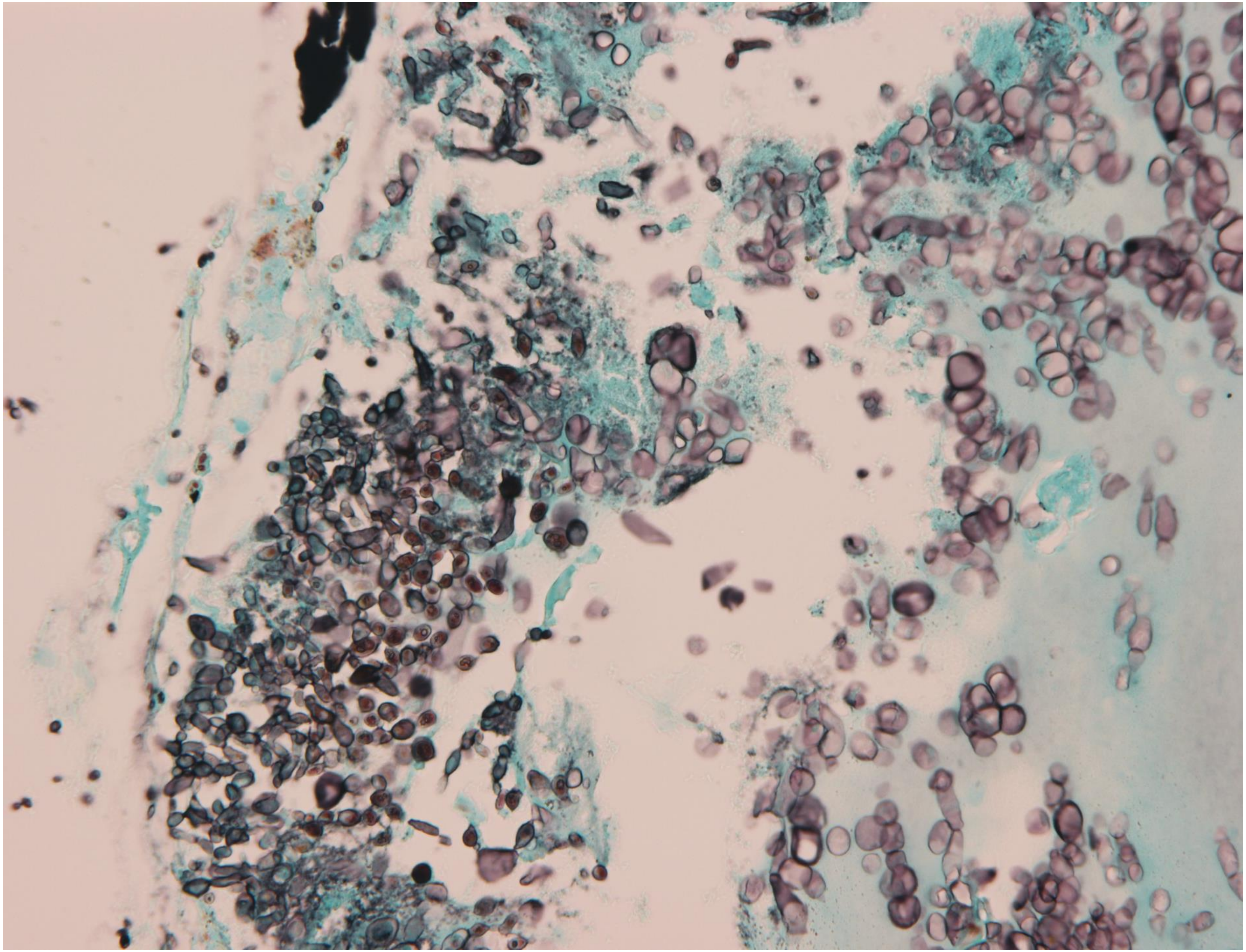

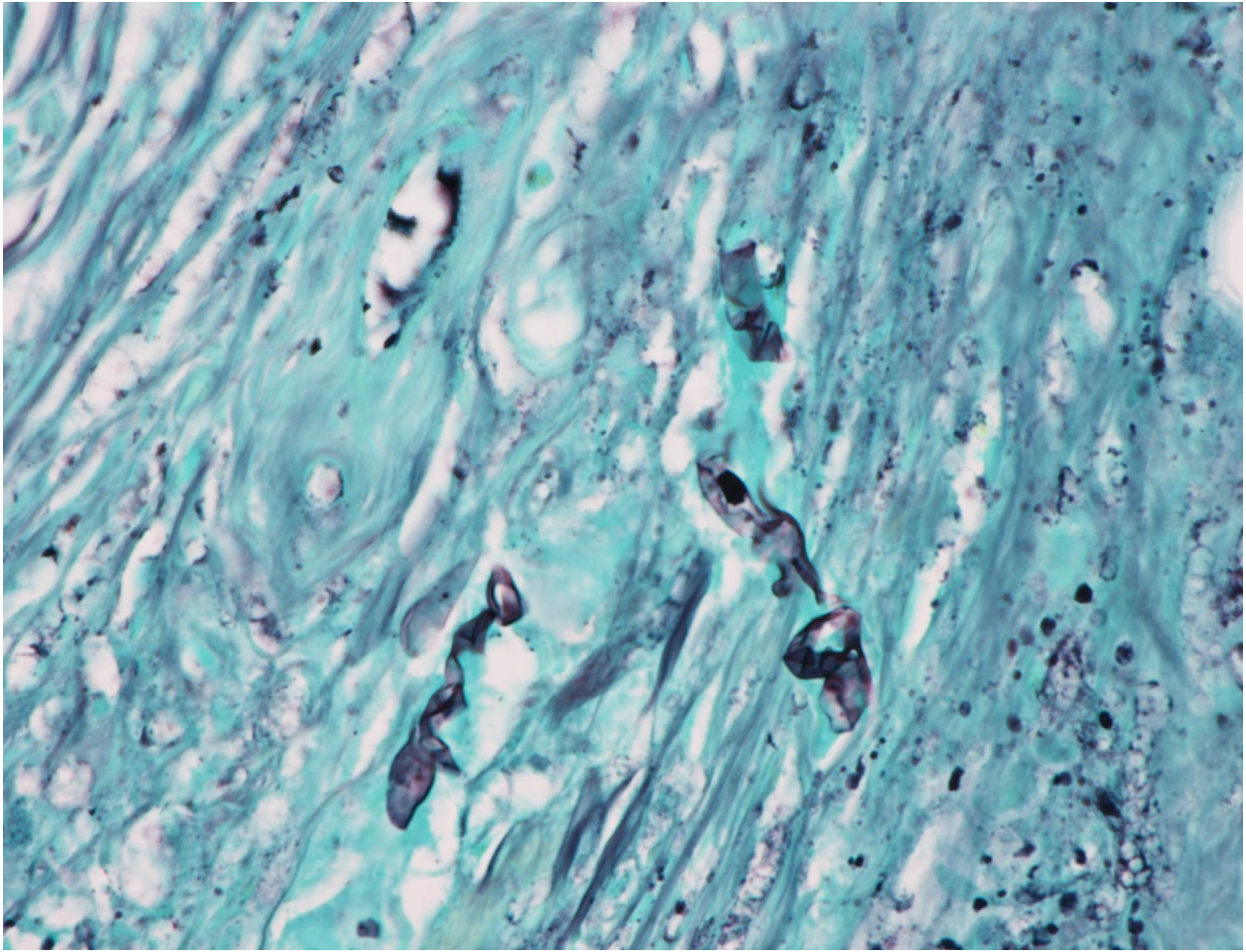

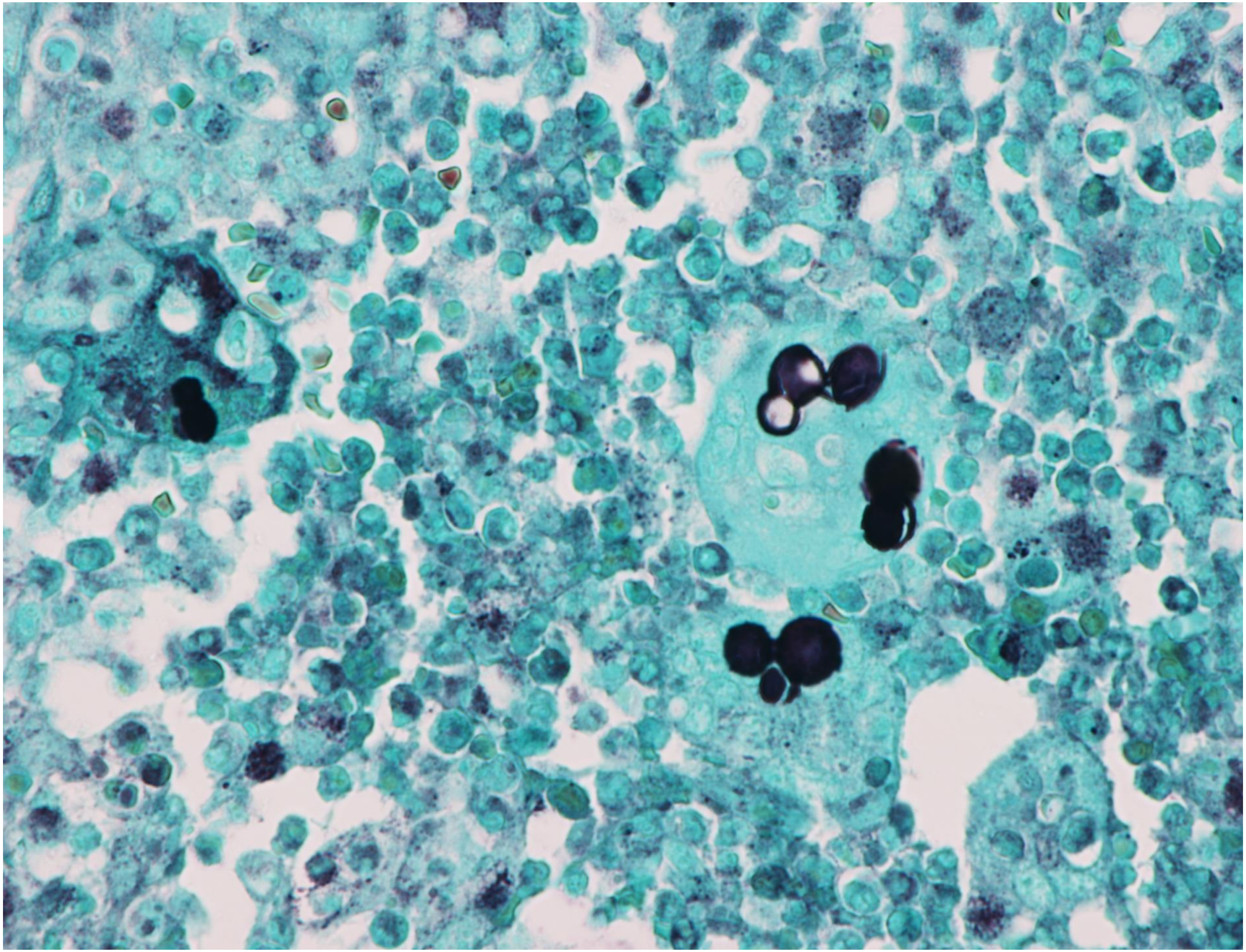

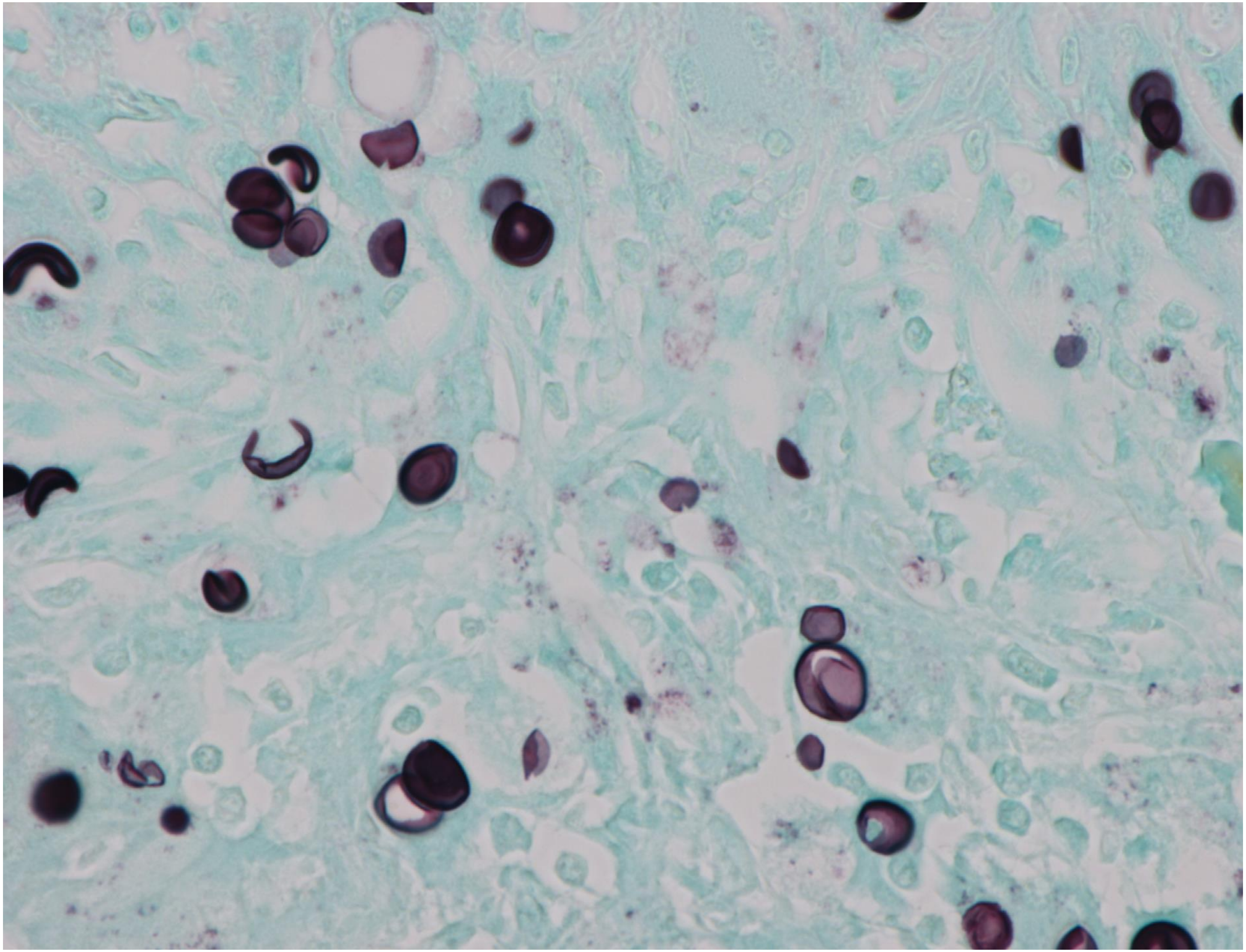

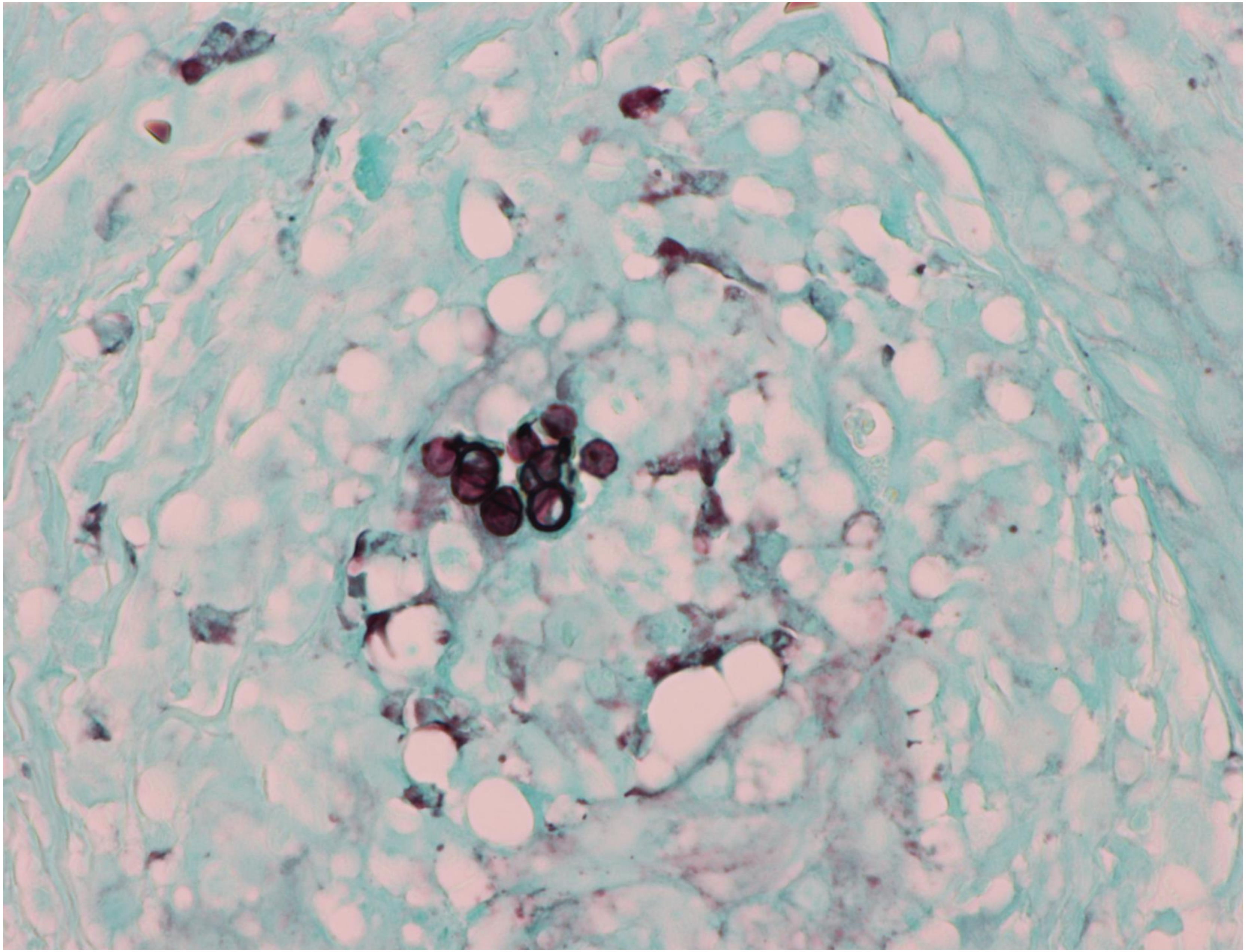

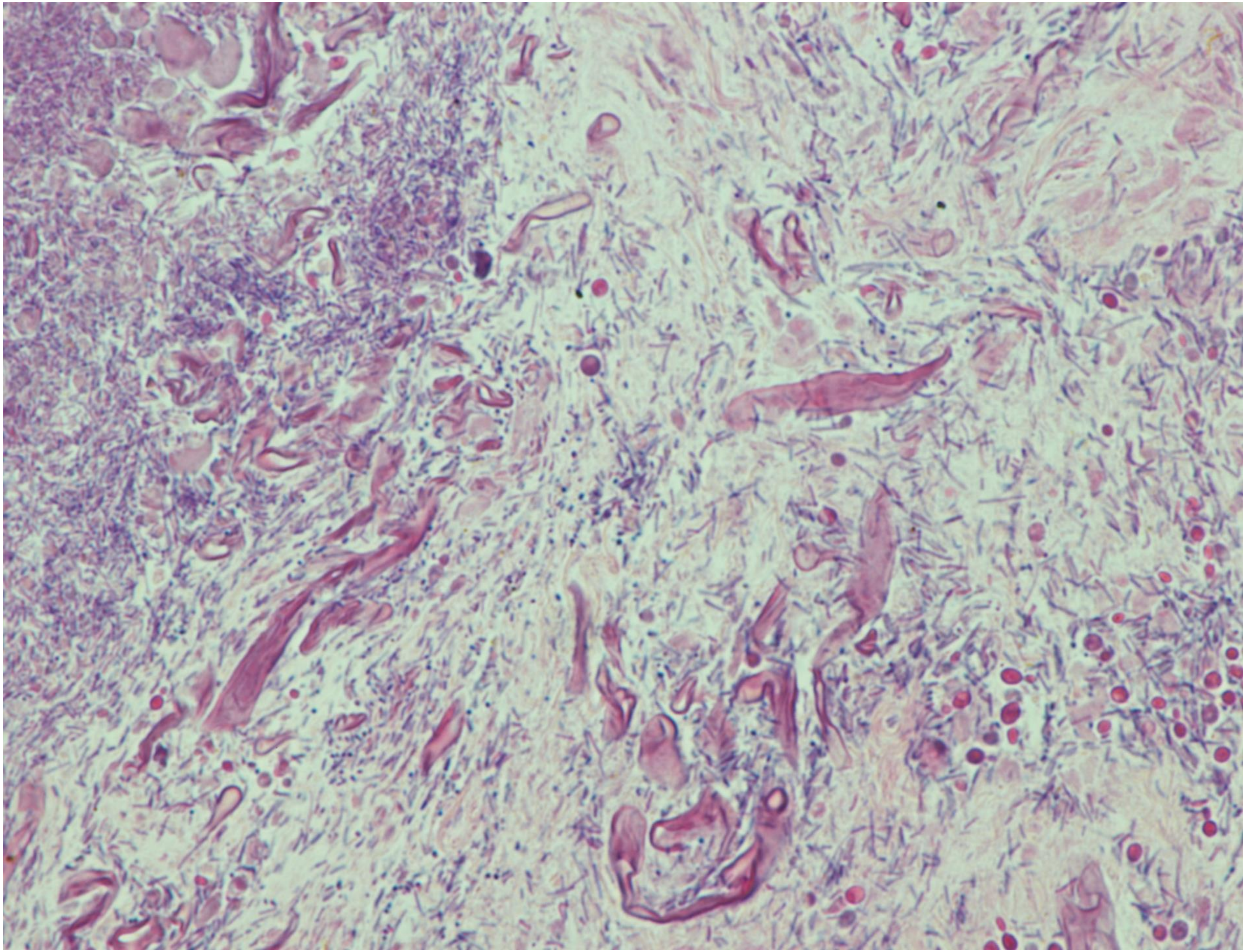

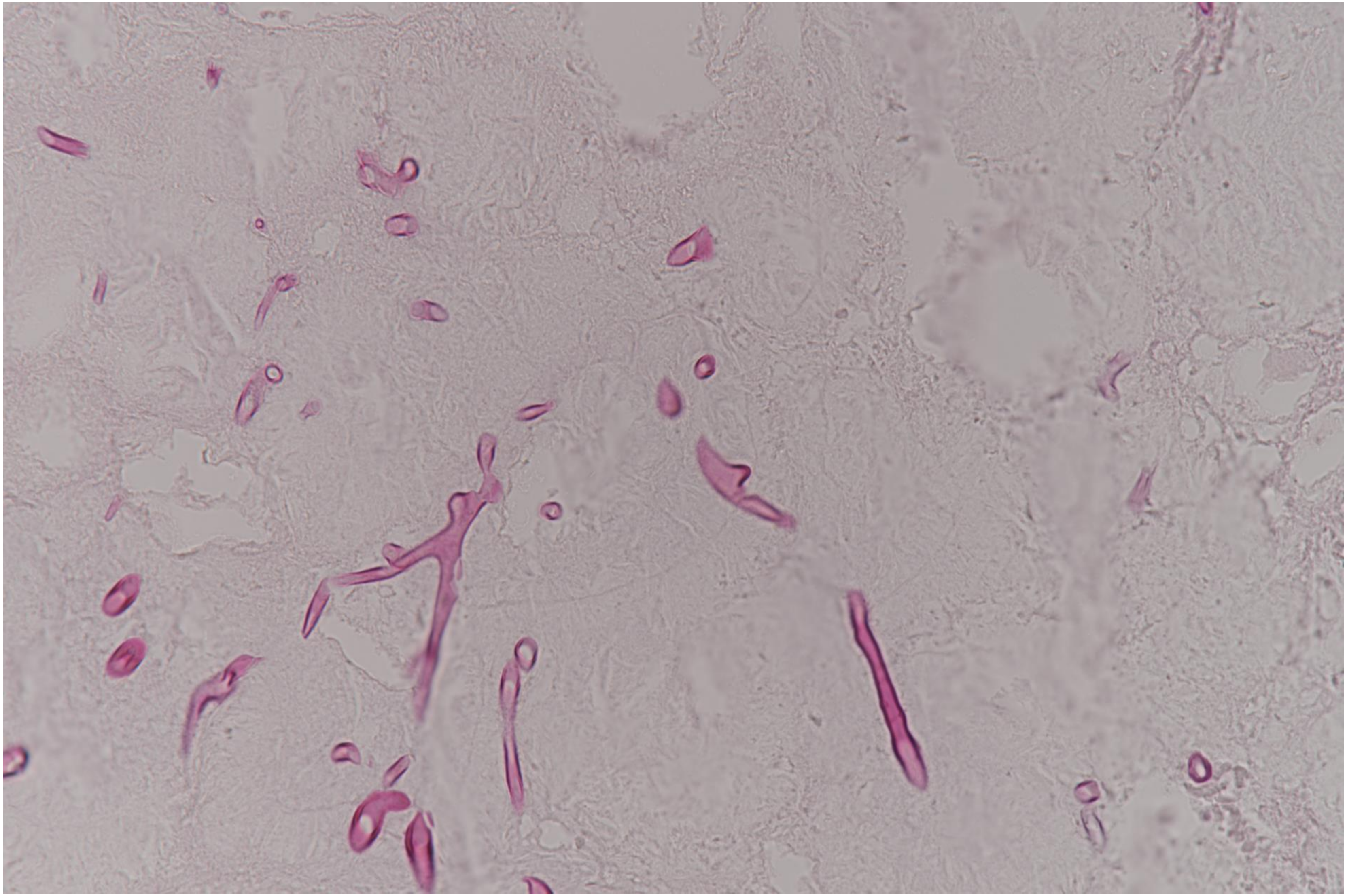

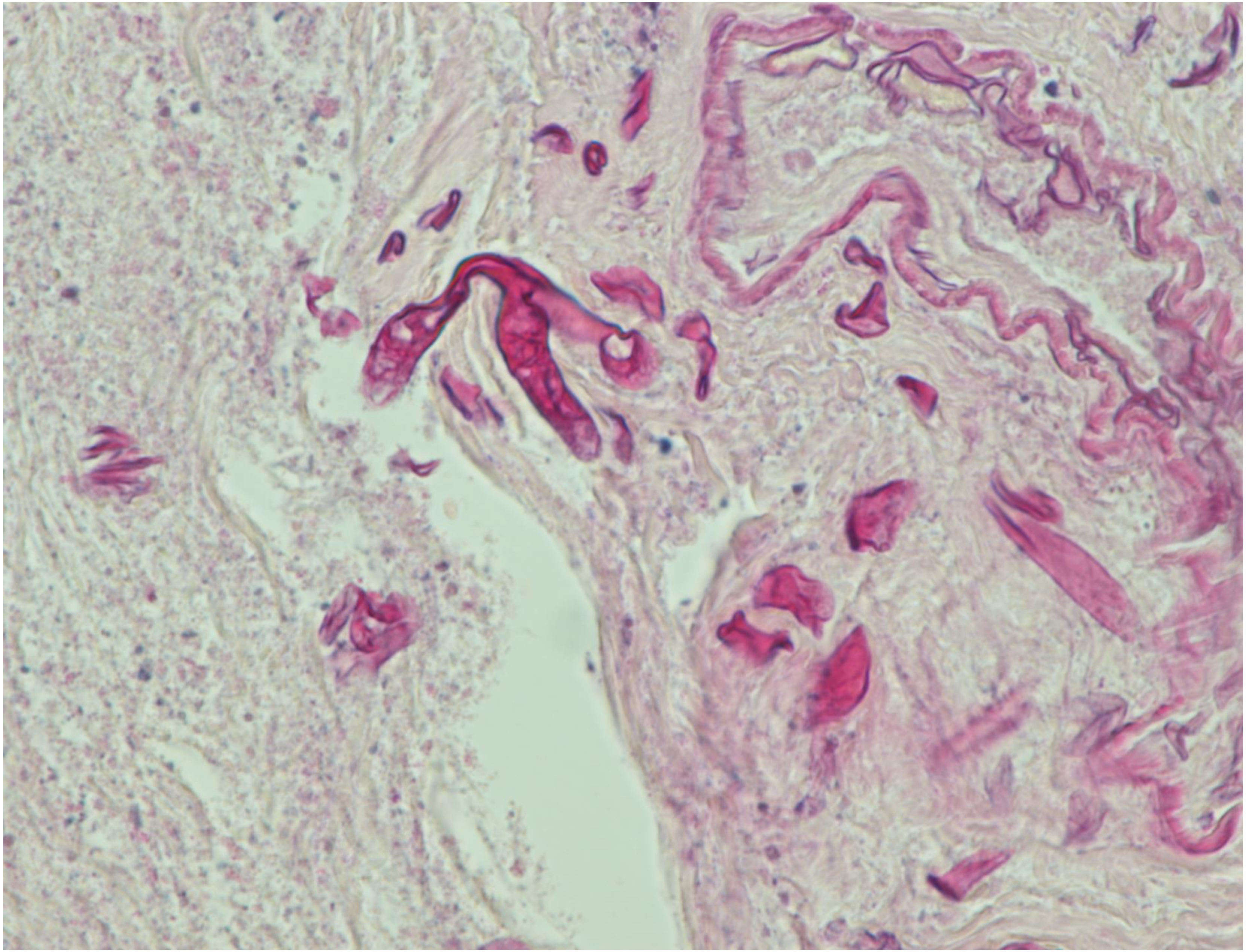

3-07

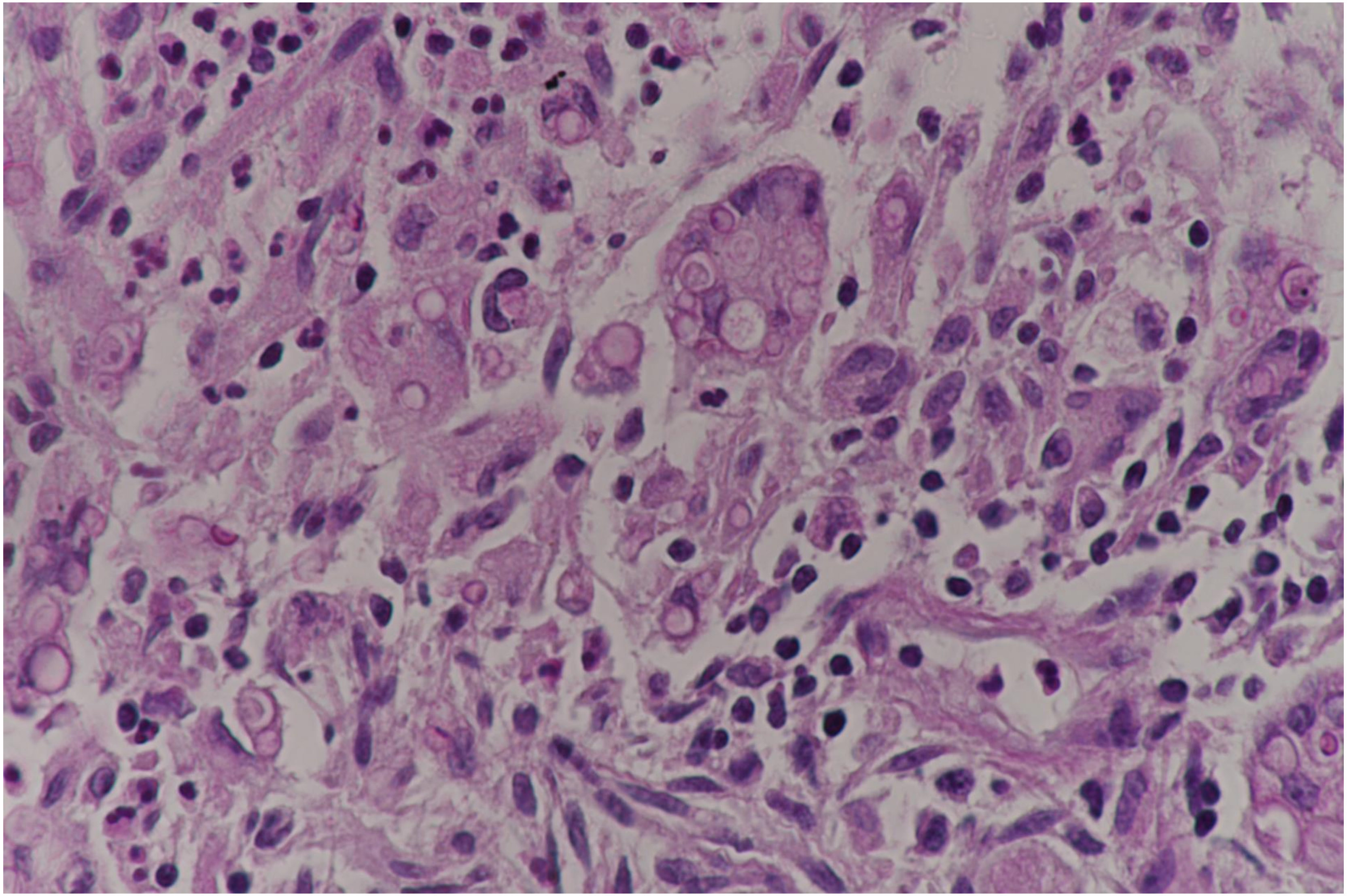

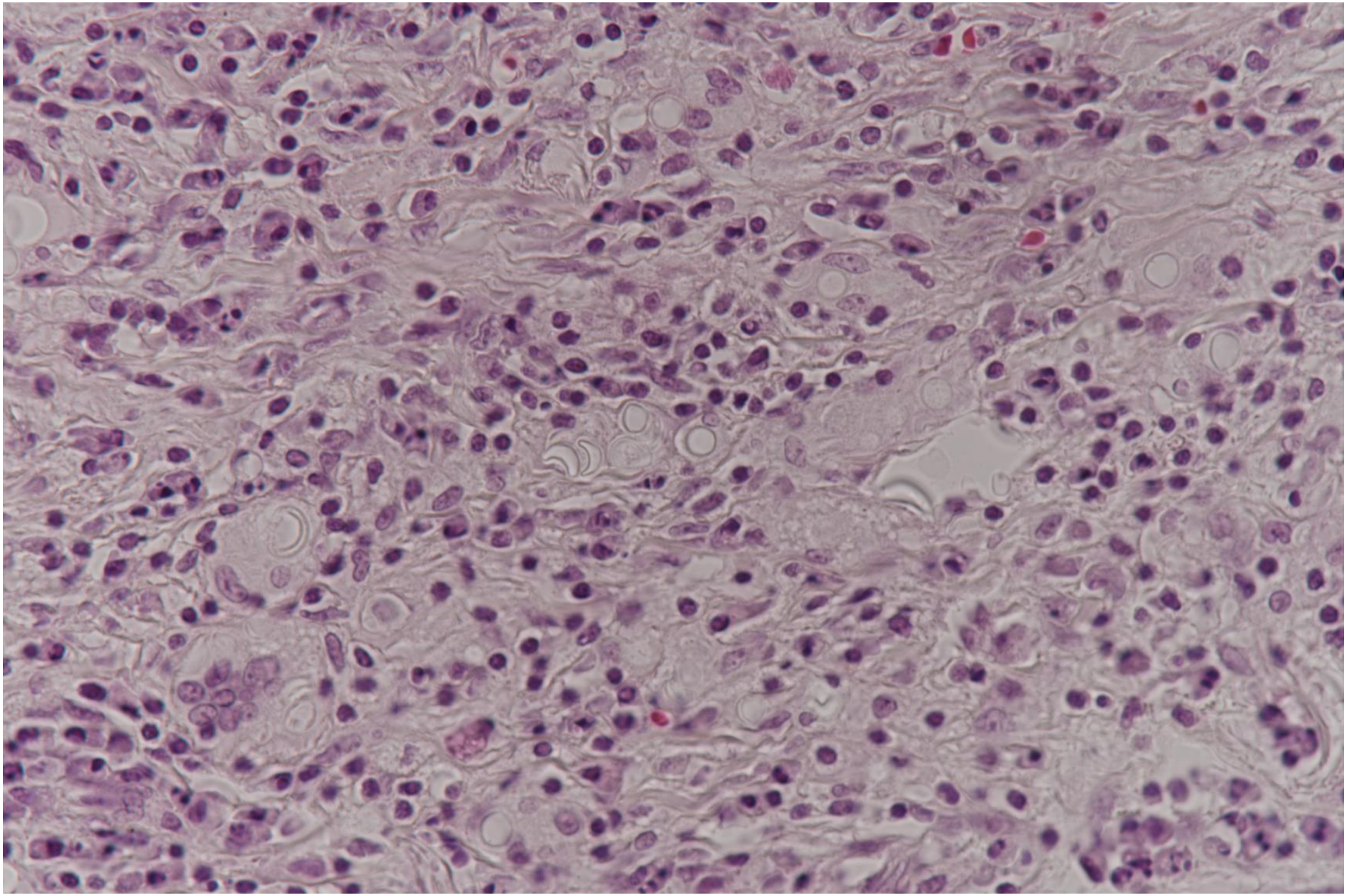

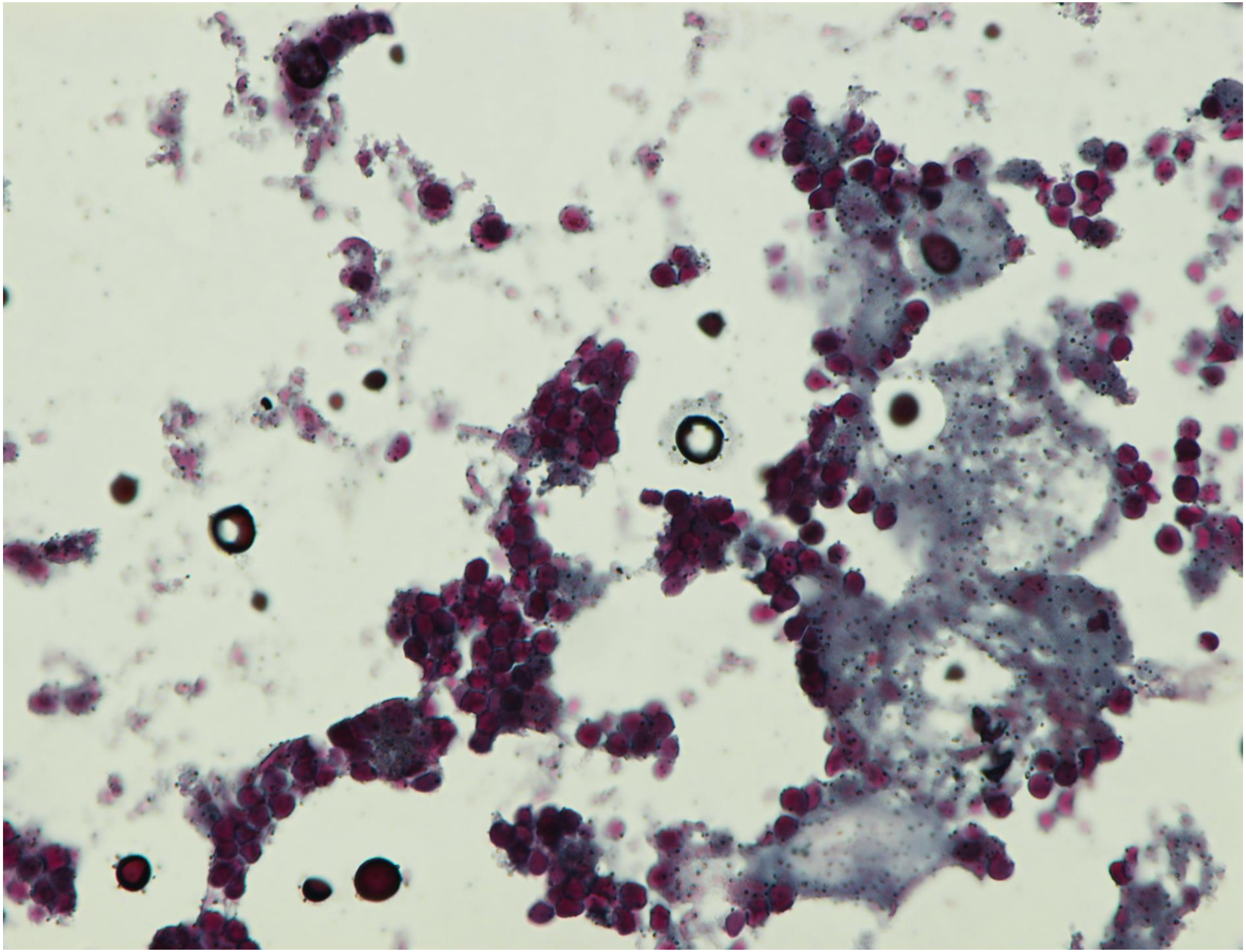

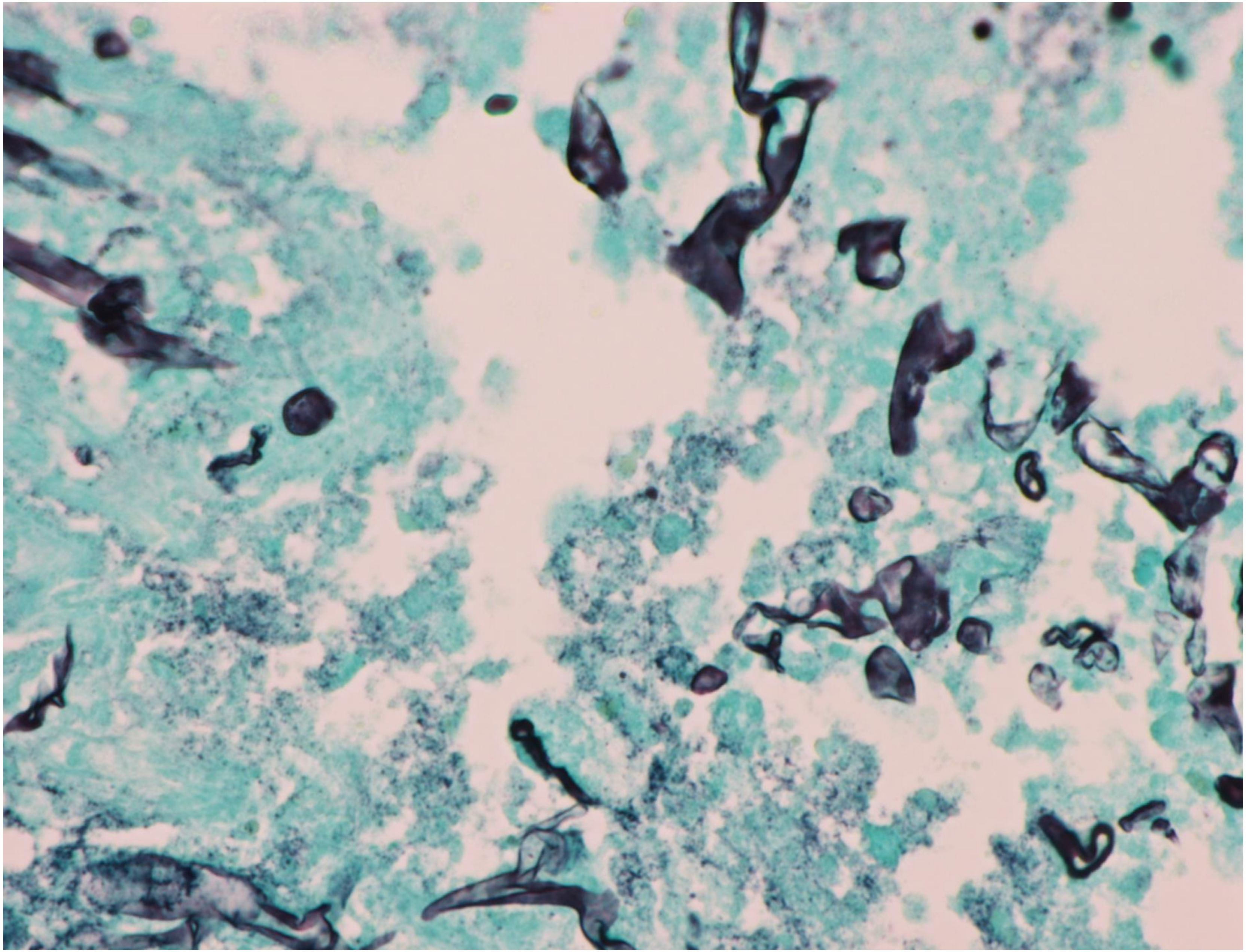

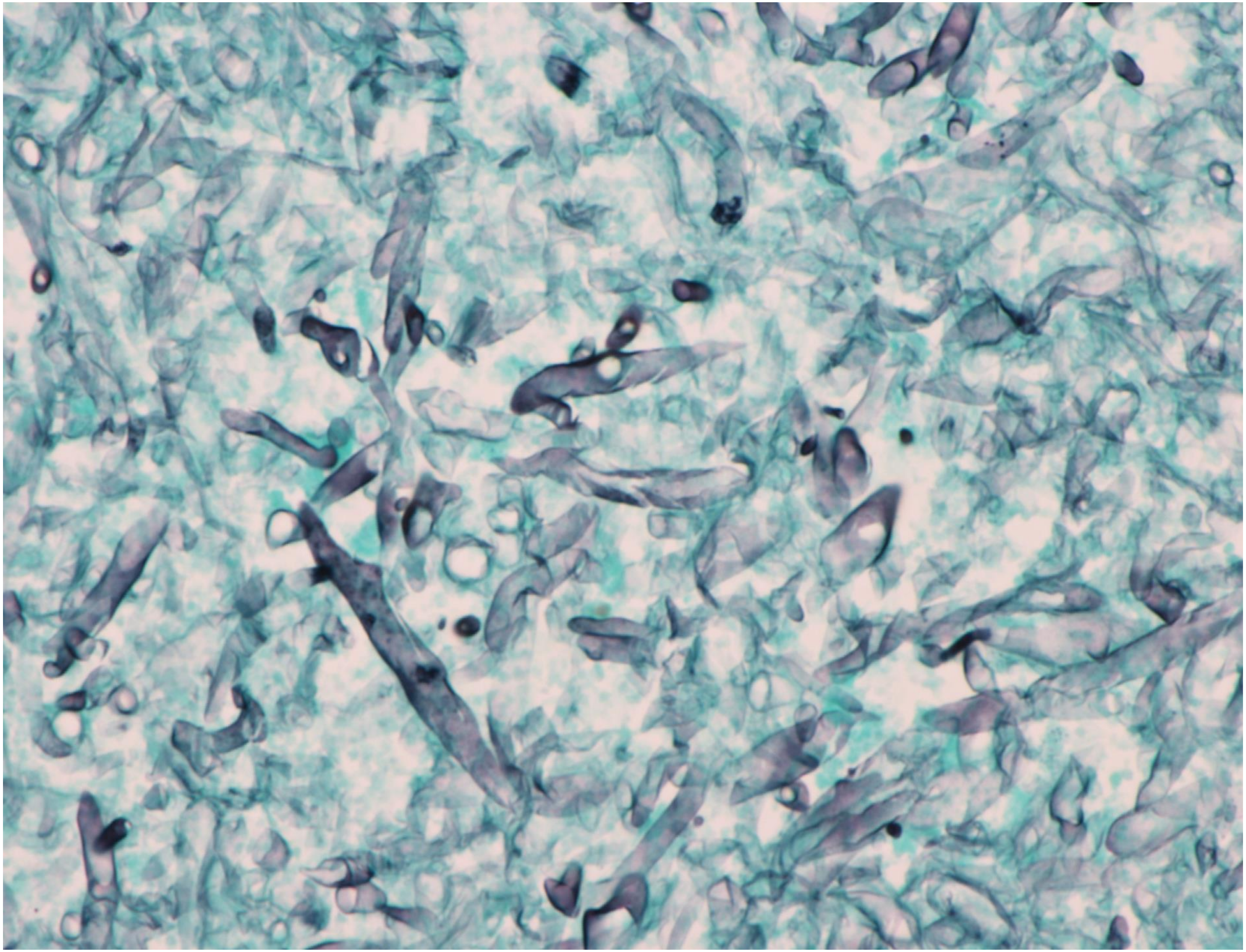

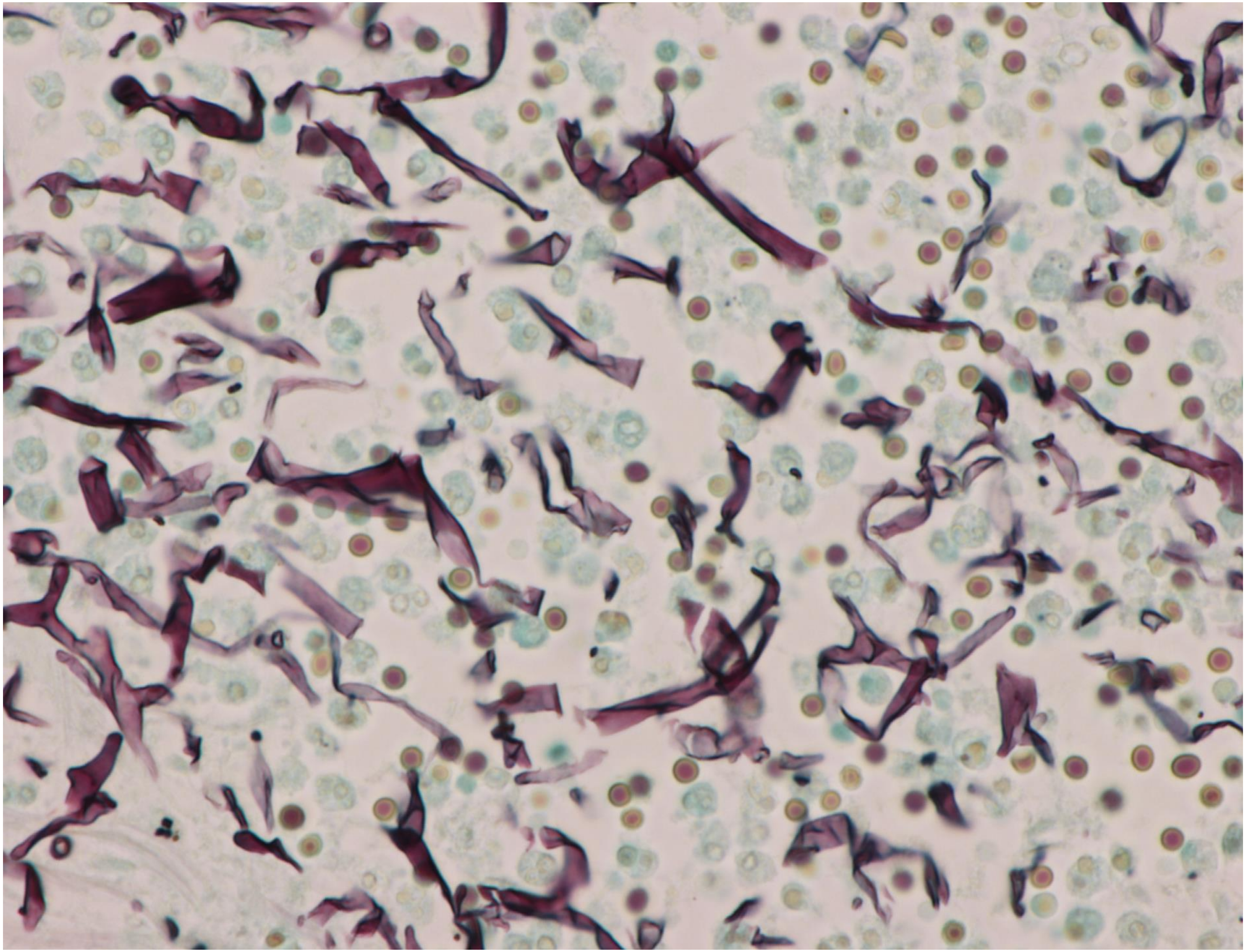

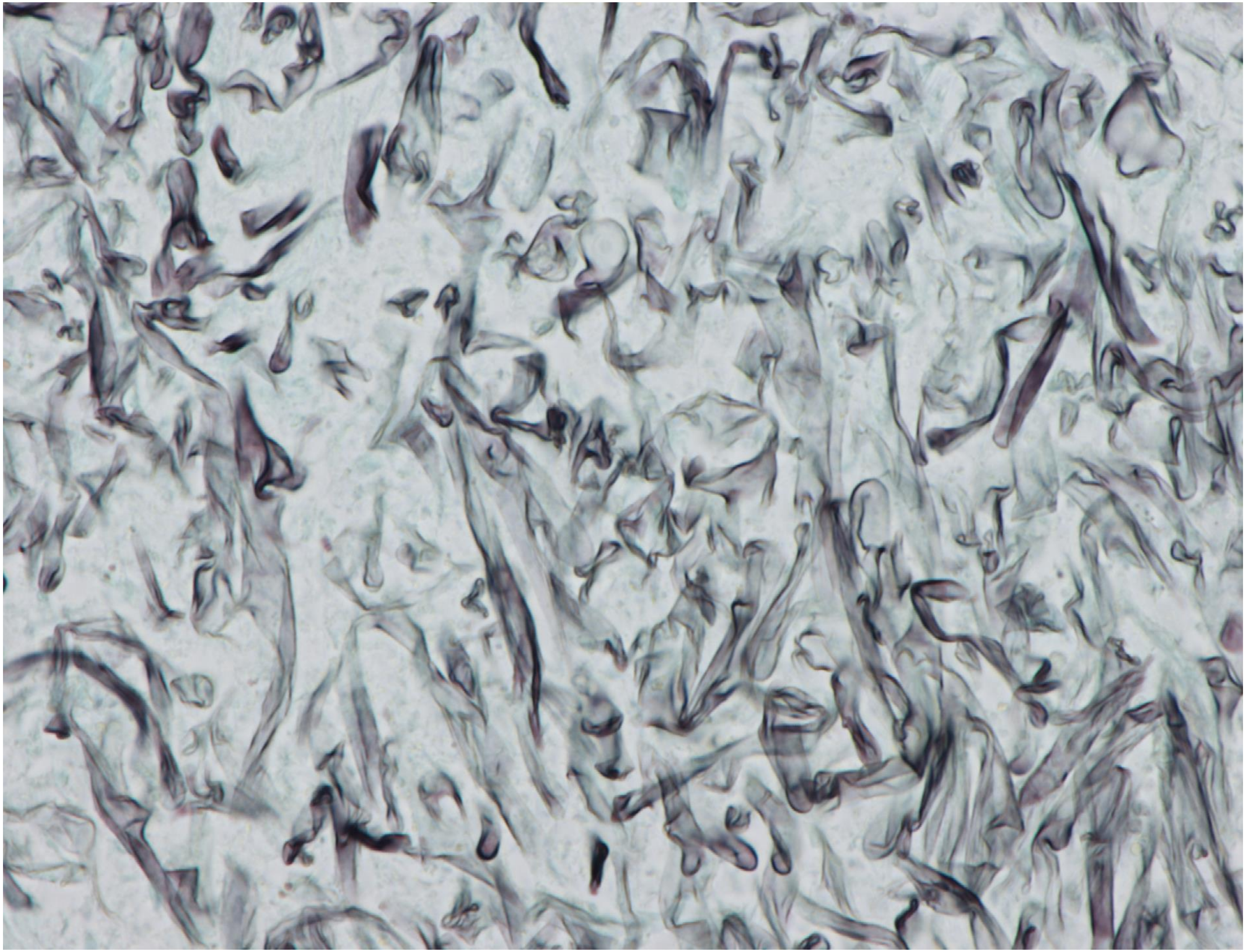

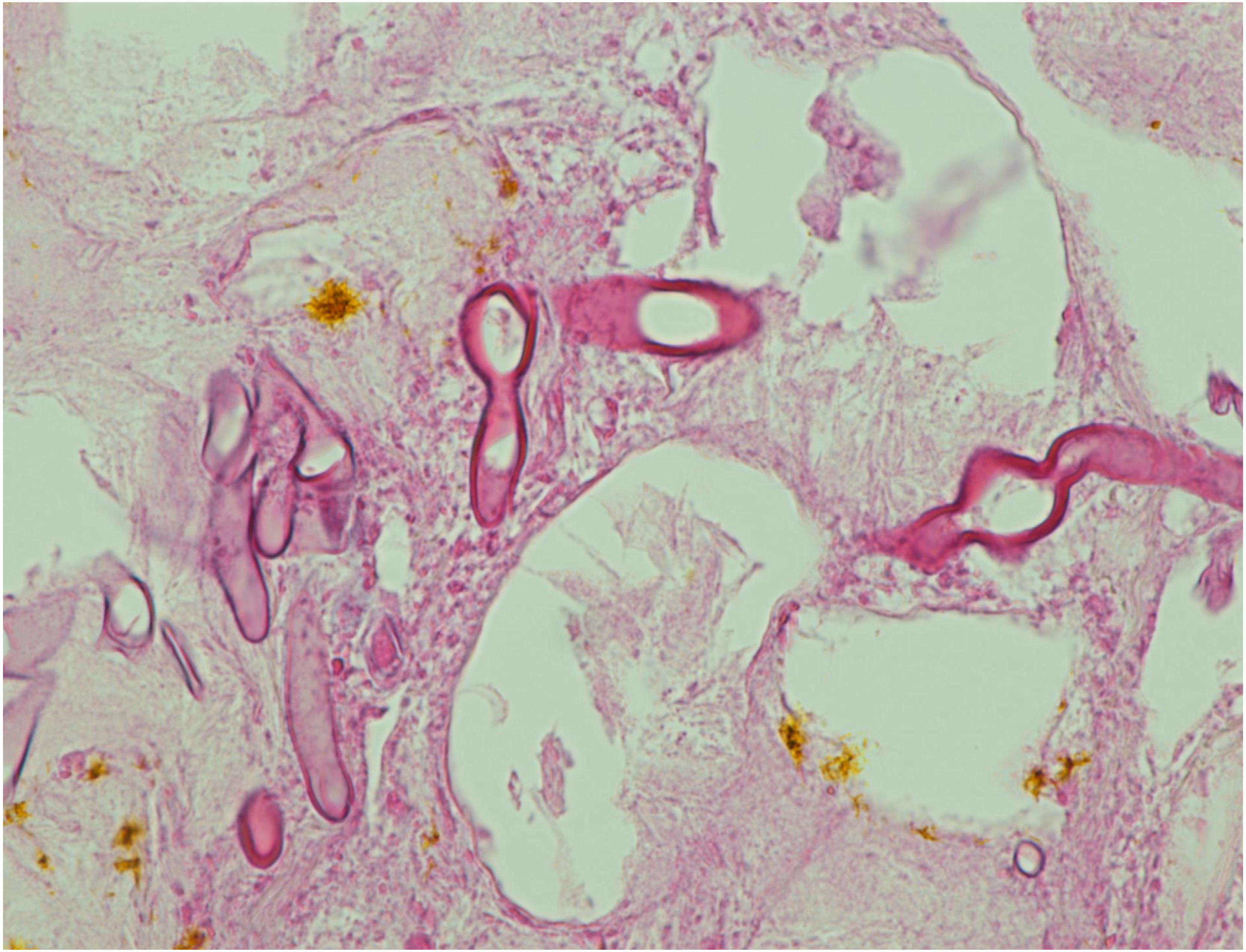

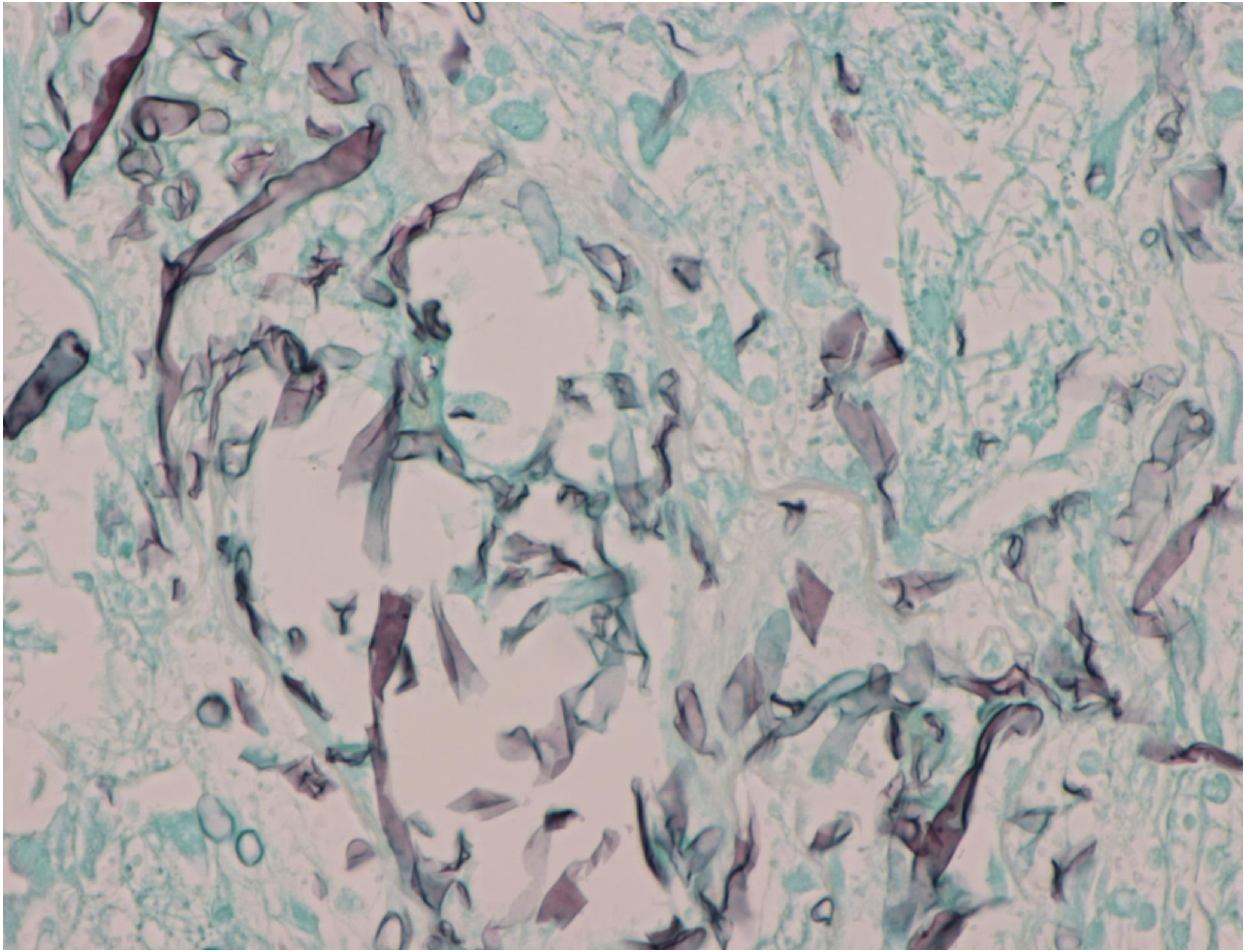

4-04

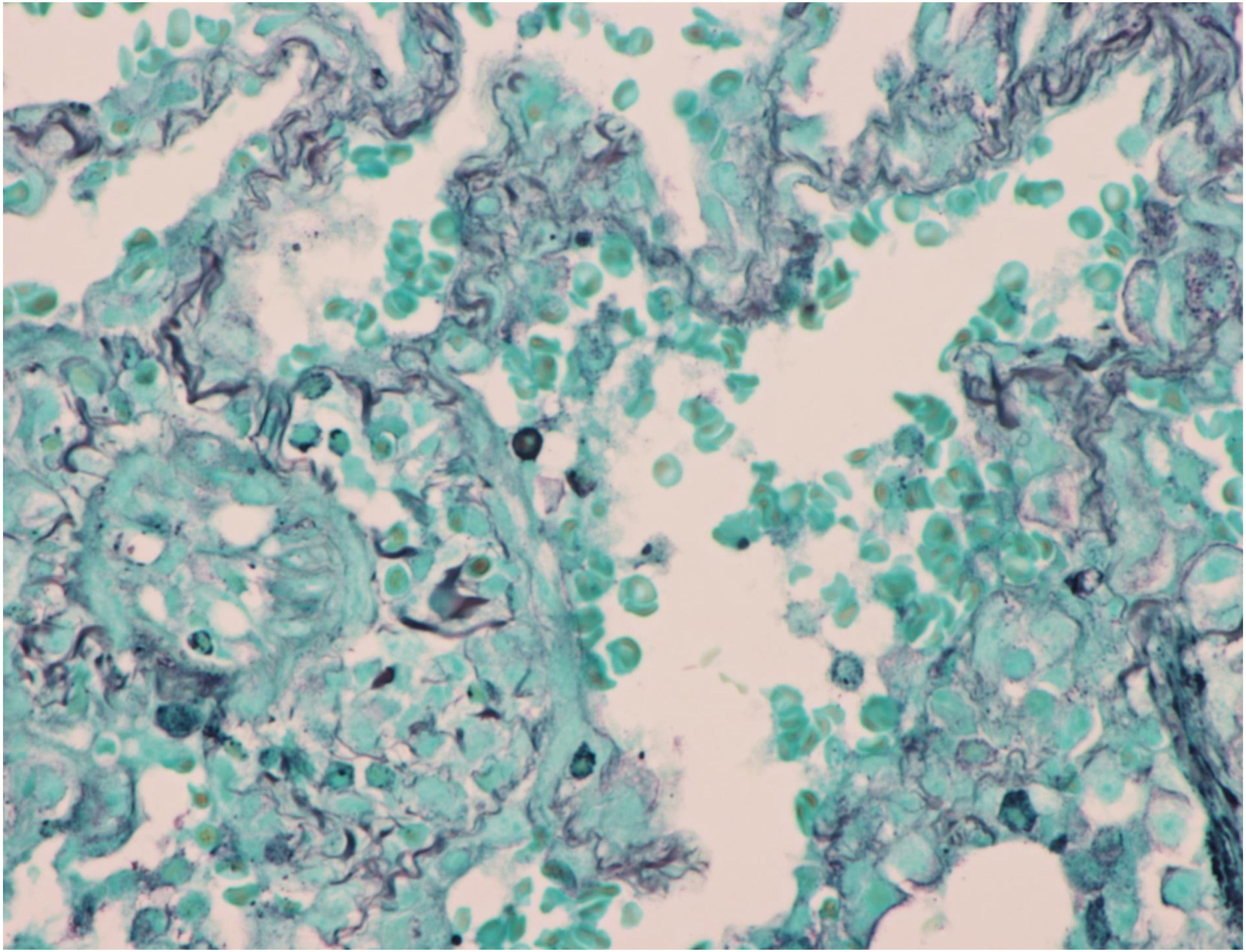

4-06

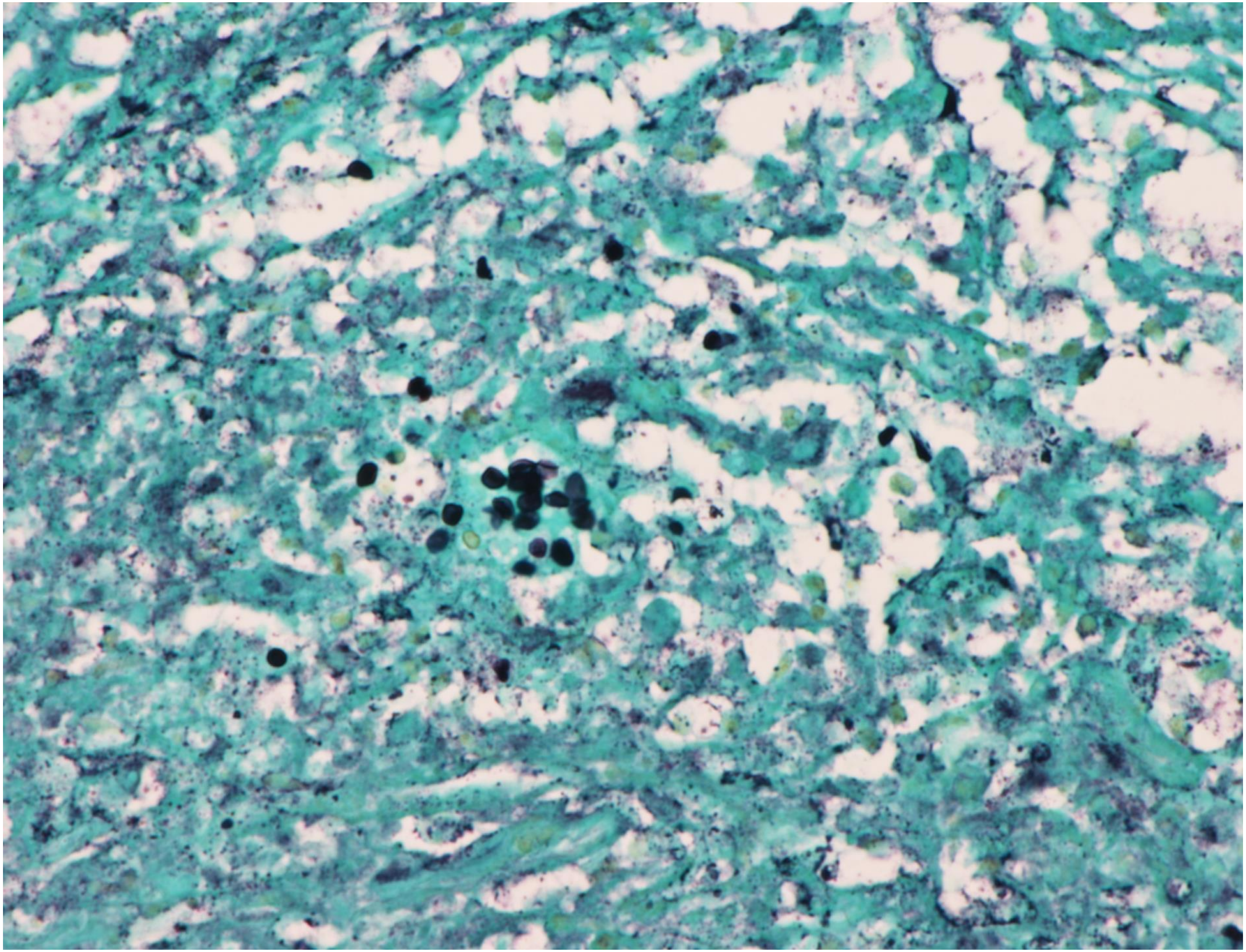

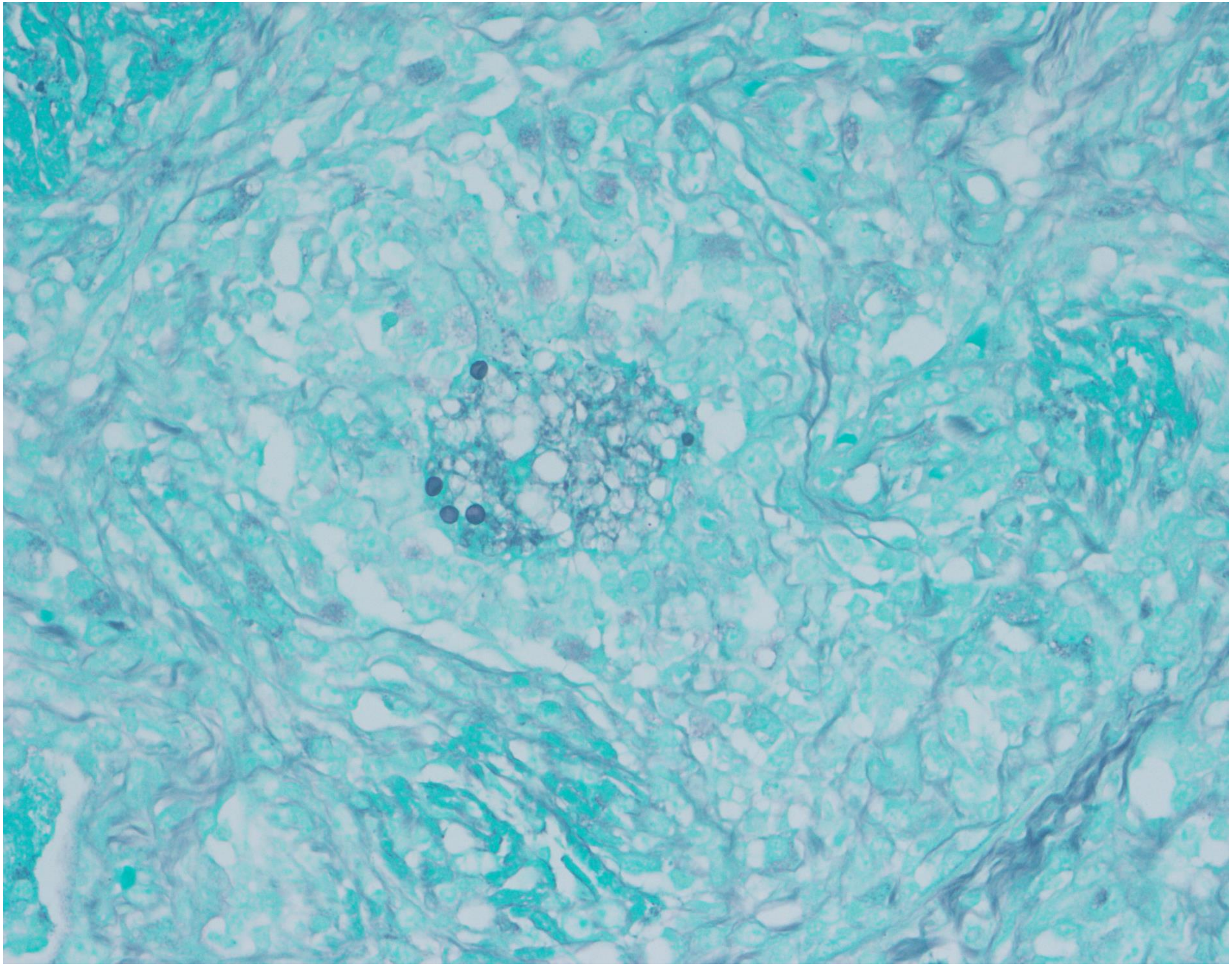

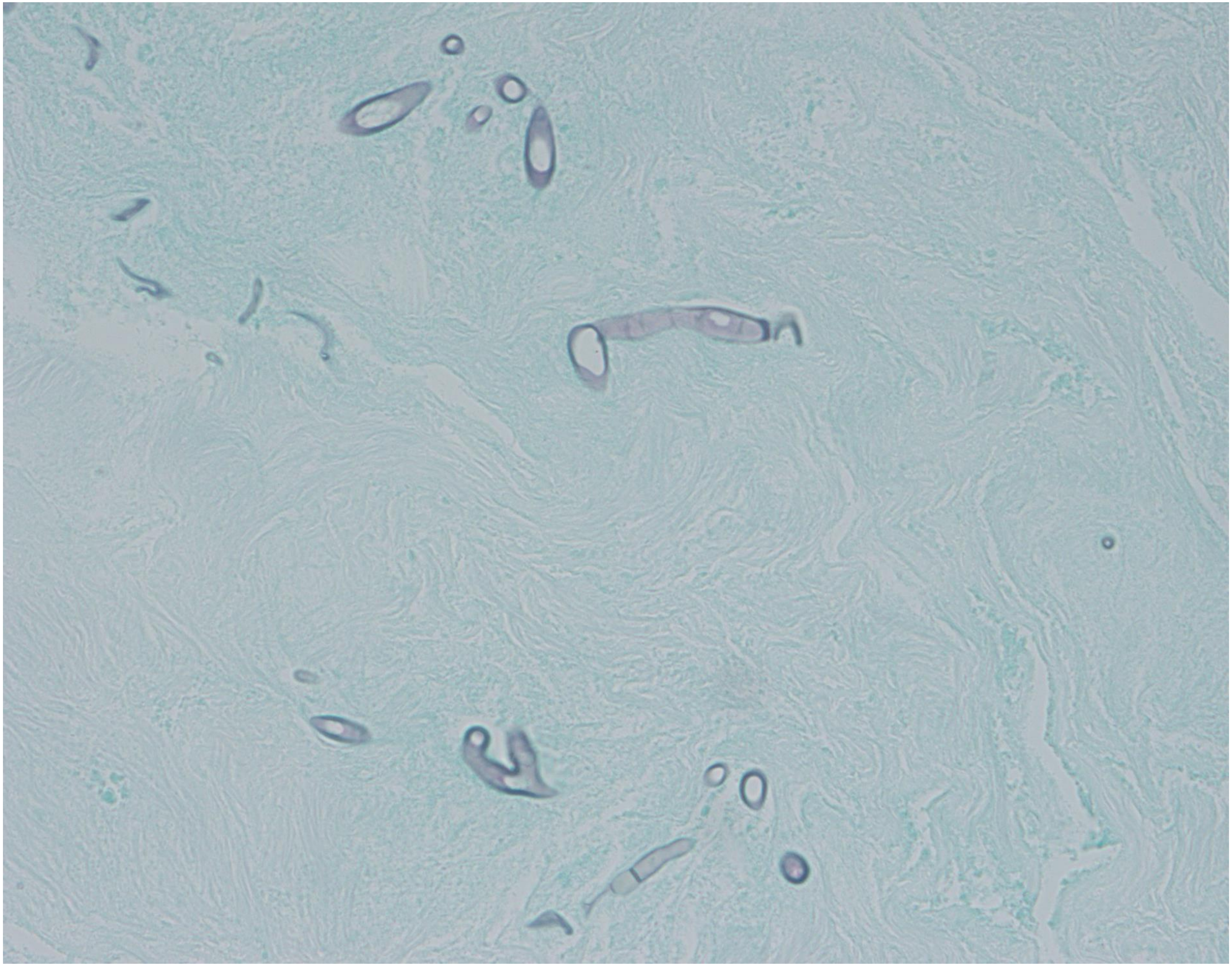

4-10

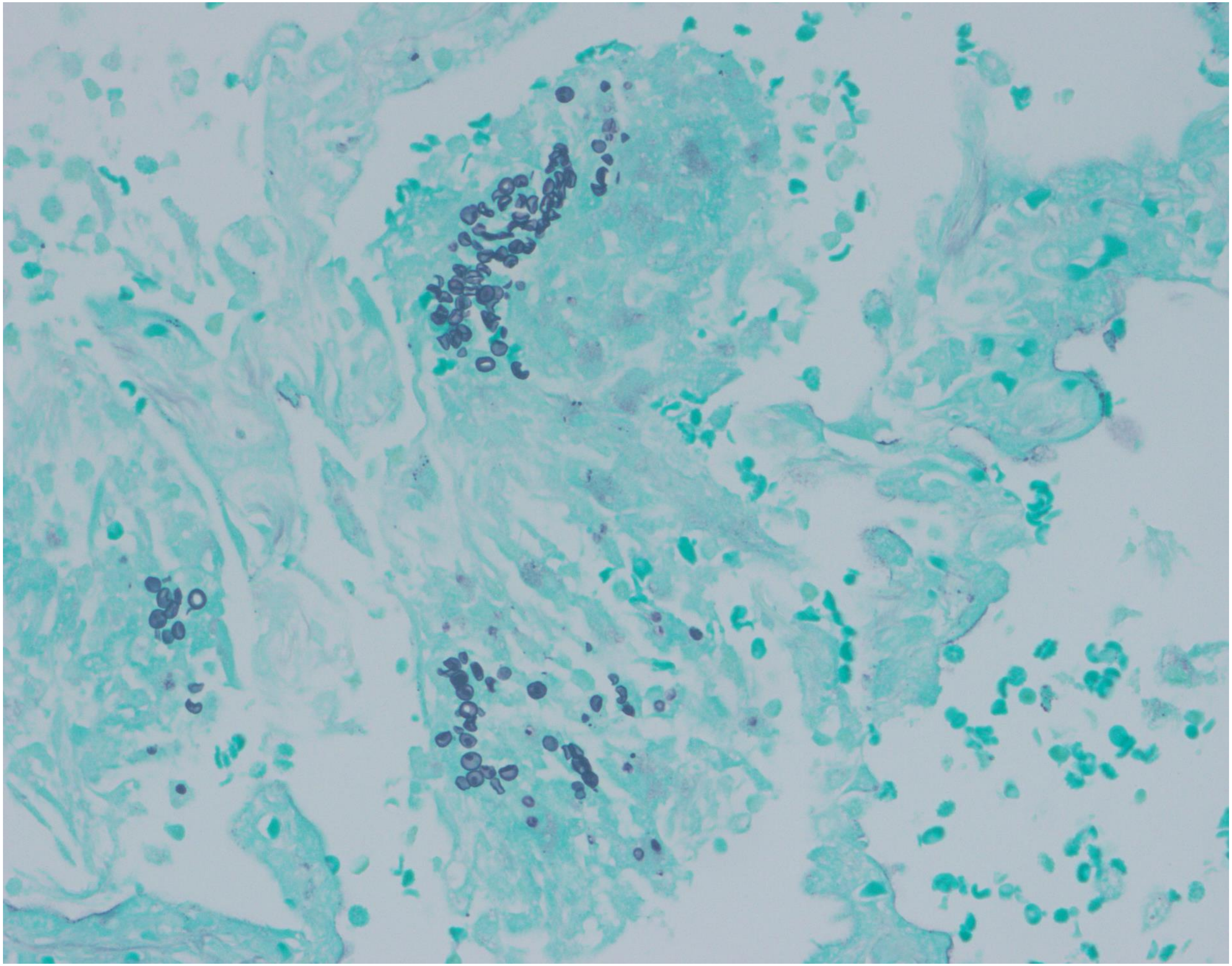

4-12

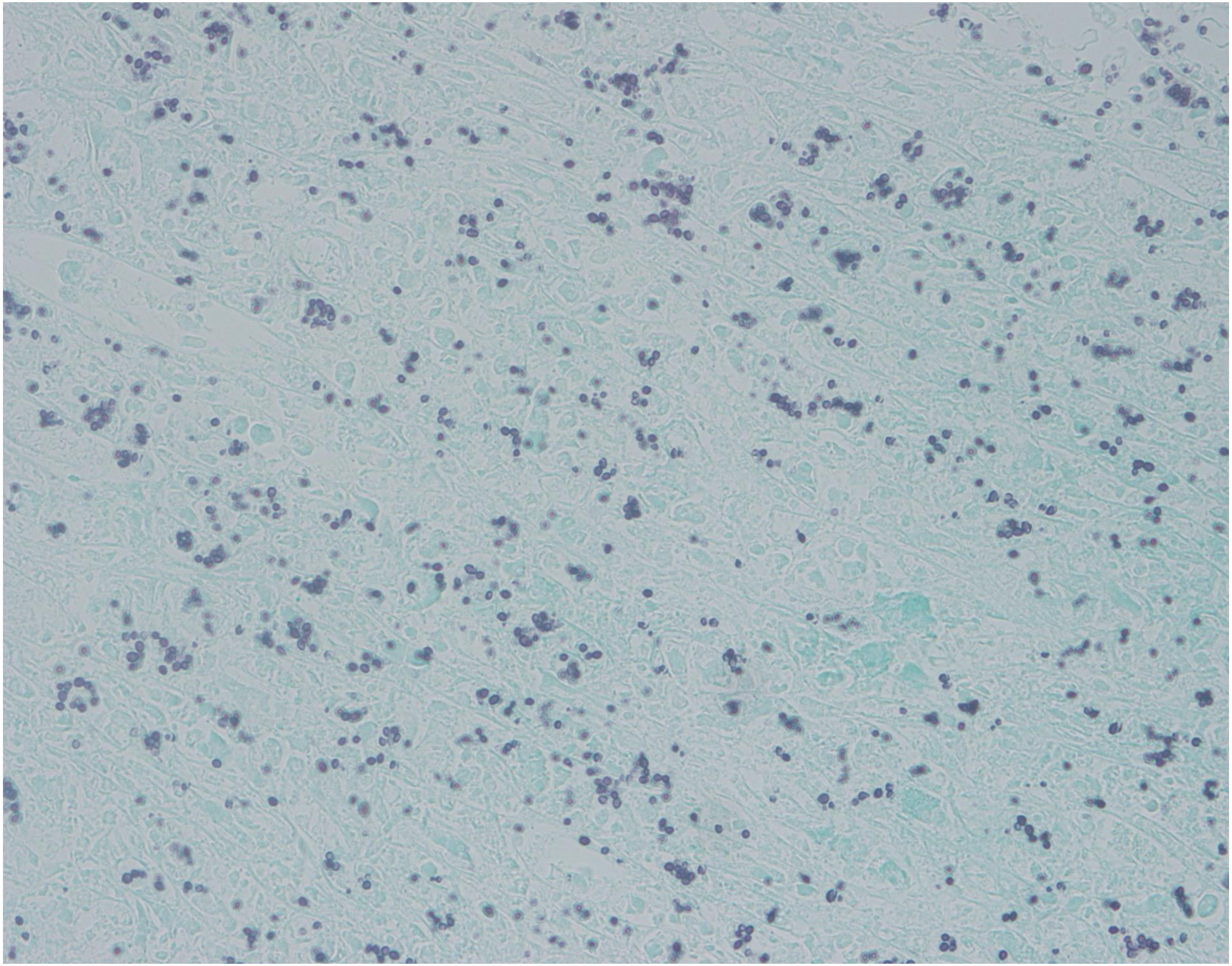

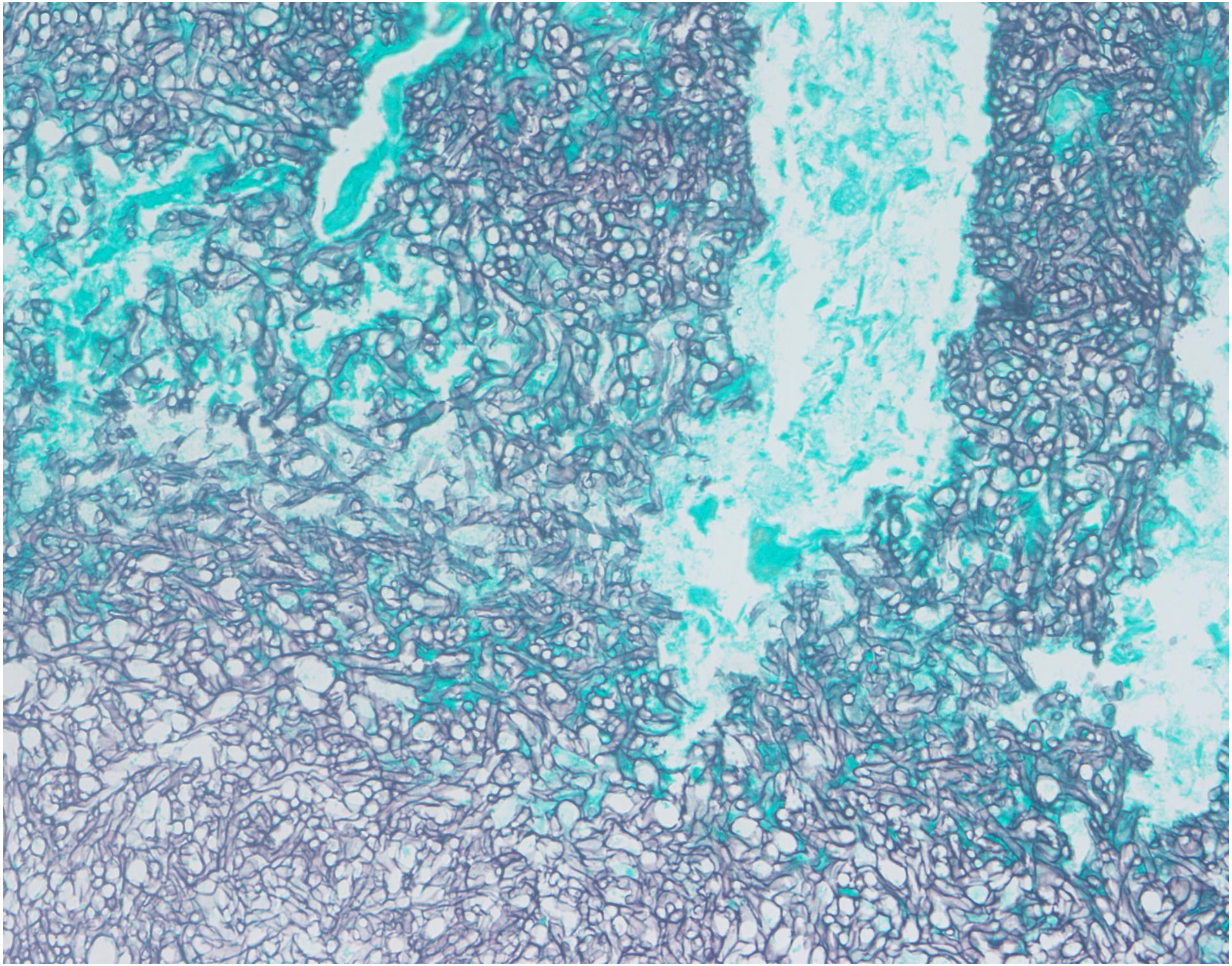

5-03

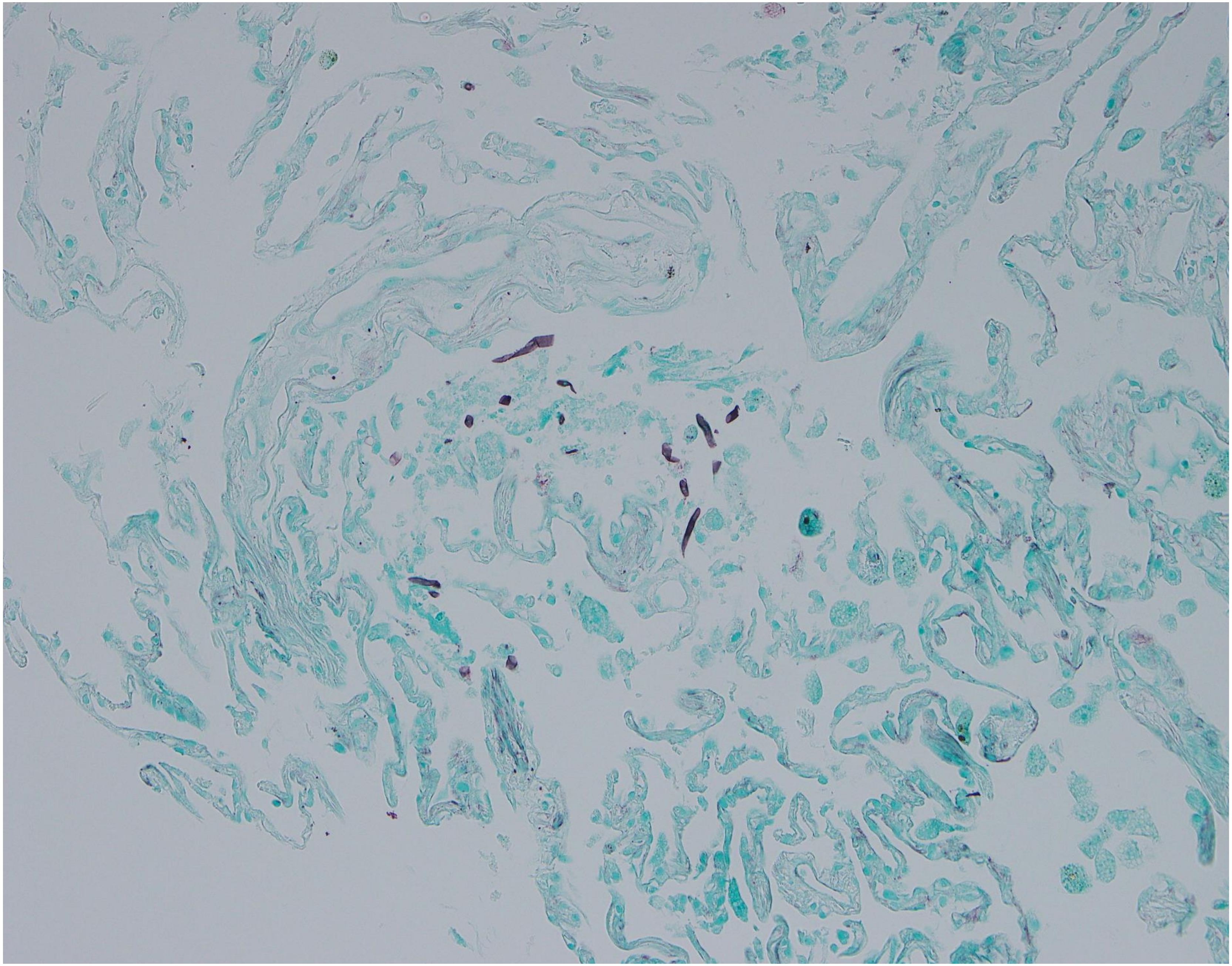

5-04

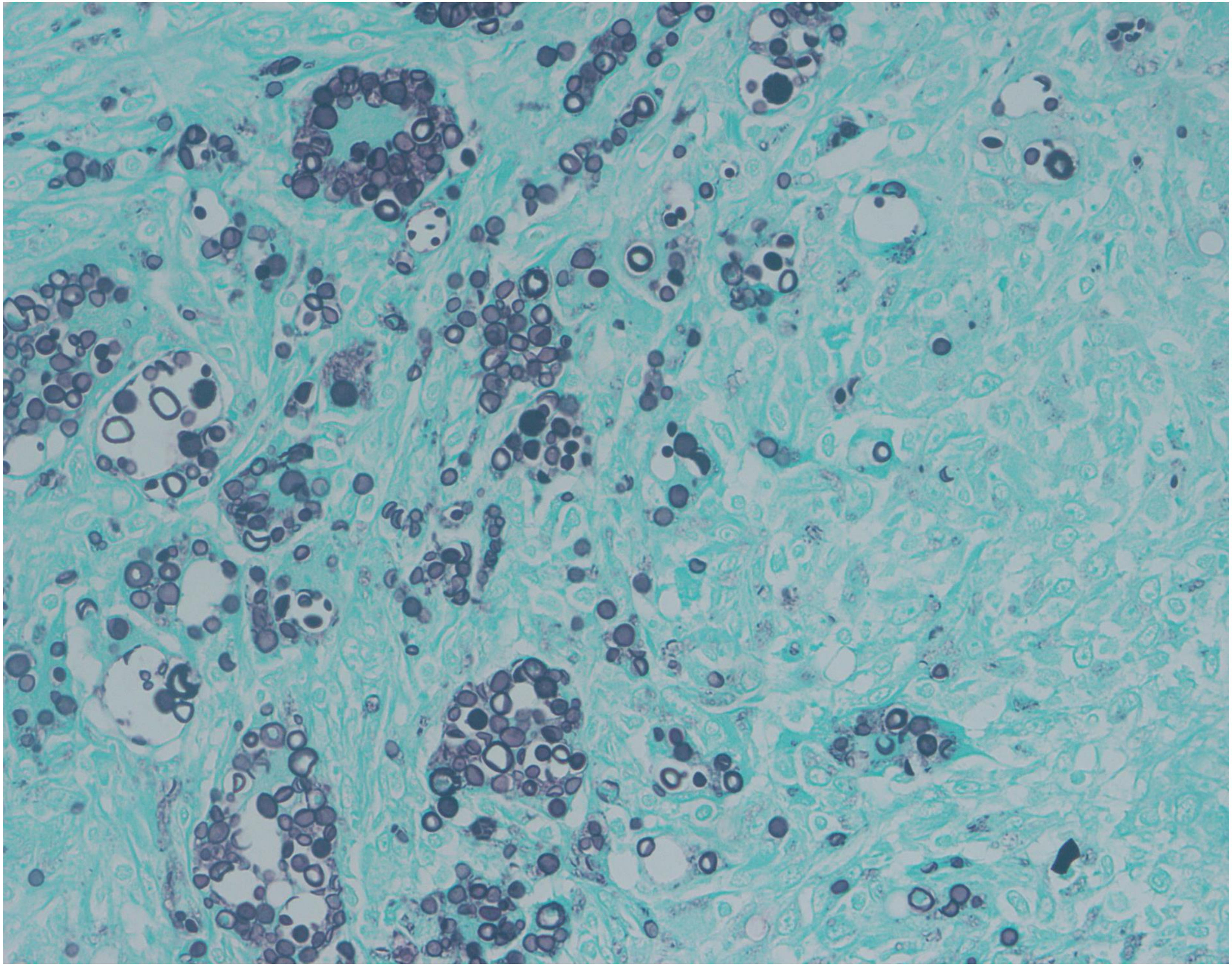

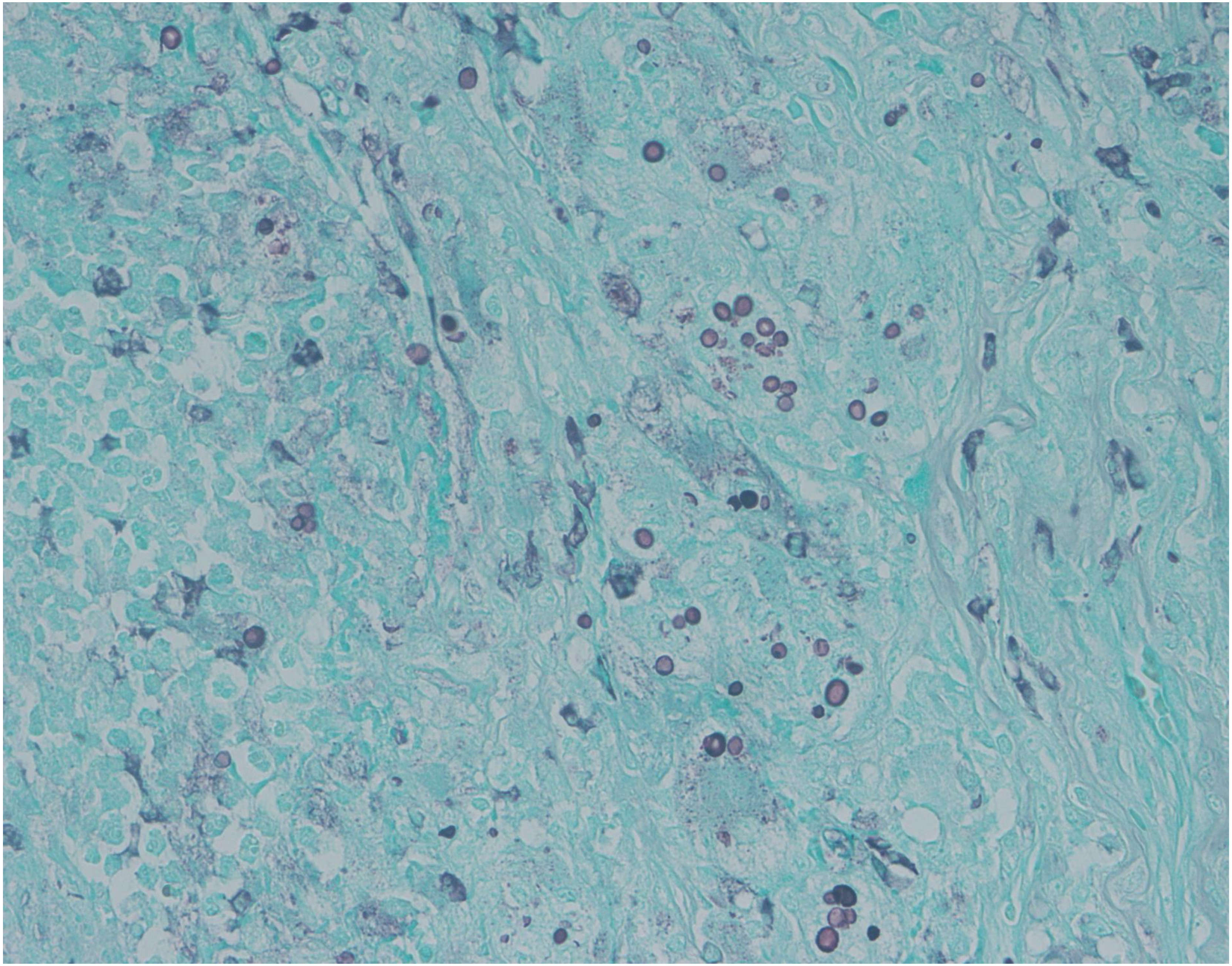

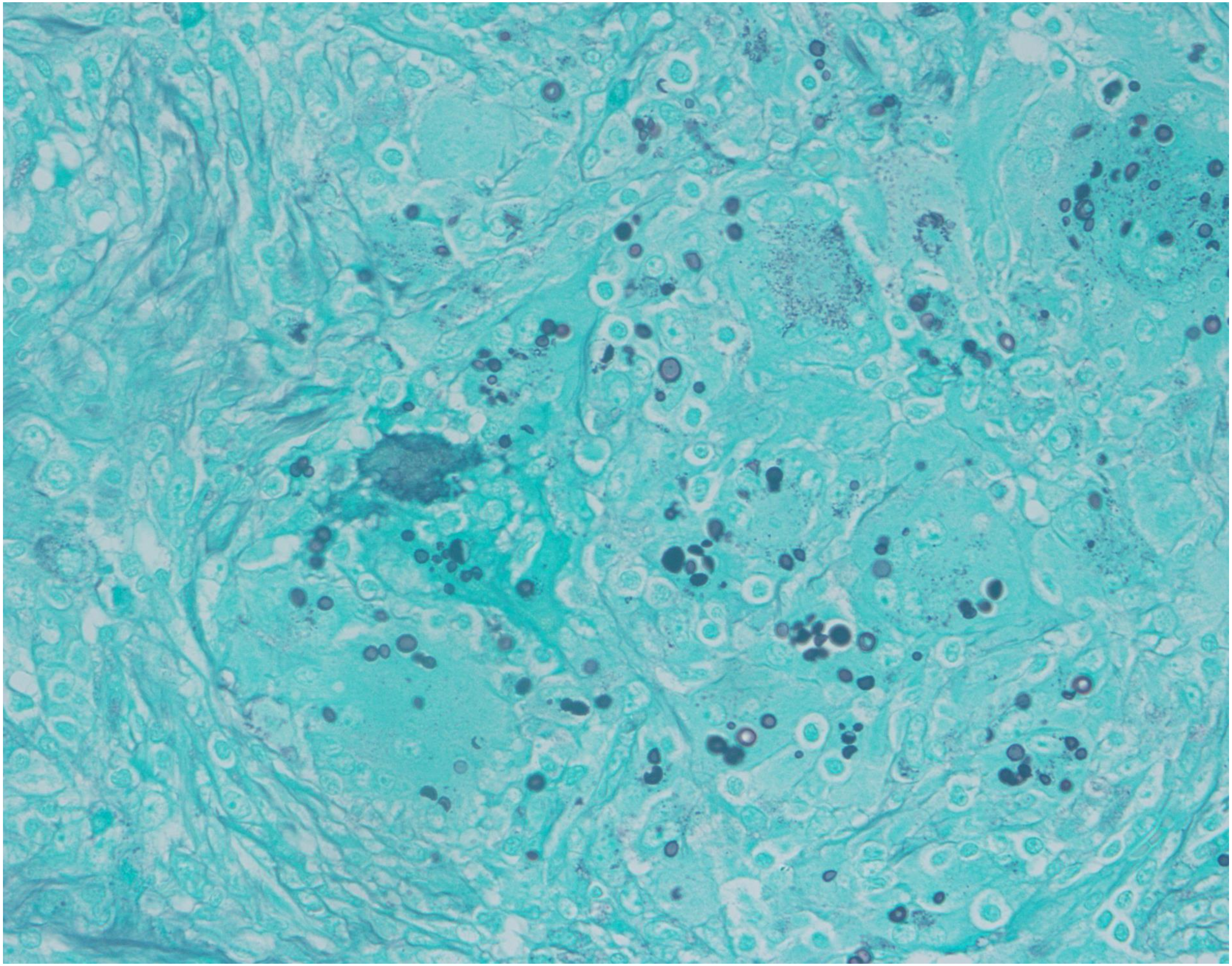

5-07
